# Mitochondrial acetyl-CoA reversibly regulates locus-specific histone acetylation and gene expression

**DOI:** 10.1101/380261

**Authors:** Oswaldo A. Lozoya, Tianyuan Wang, Dagoberto Grenet, Taylor C. Wolfgang, Mack Sobhany, Douglas Ganini da Silva, Gonzalo Riadi, Navdeep Chandel, Richard P. Woychik, Janine H. Santos

## Abstract

The impact of mitochondria in epigenetics is emerging but our understanding of this relationship and its impact on gene expression remain incomplete. We previously showed that acute mitochondrial DNA (mtDNA) loss leads to histone hypoacetylation. It remains to be defined if these changes are maintained when mitochondrial dysfunction is chronic and, importantly, if they are sufficient to alter gene expression. To fill these gaps, we here studied both a progressive and a chronic model of mtDNA depletion using biochemical, pharmacological, genomics and genetic assays. We show that histones are hypoacetylated in both models. We link these effects to decreased histone acetyltransferase (HAT) activity independent of changes in ATP citrate lyase function, which can be reversibly modulated by altering specifically the mitochondrial pool of acetyl-CoA. Also, we determined that these changes regulate locus-specific gene expression and physiological outcomes, including the production of prostaglandins. These results may be relevant to the pathophysiology of mtDNA depletion syndromes and to understanding the effects of environmental agents, such as AZT or antibiotics, that lead to physical or functional mtDNA loss.

## Introduction

The role of mitochondria in cell biology and organismal health has expanded dramatically in the last decade. From a focus originally on bioenergetics, it is now recognized that mitochondria broadly affect cell physiology in diverse ways. For instance, mitochondria interact with other organelles, such as the endoplasmic reticulum, by close contacts or through the generation of small vesicular carriers, which allows the transport and exchange of lipids, proteins and other small molecules like calcium [1,2]. Mitochondria are also important players in signaling via reactive oxygen species (ROS) and other metabolites that impart post-translational modifications to many proteins, including transcription factors [3]. Most recently, we and others have shown that mitochondria influence the epigenome [4-8], yet full mechanistic insights and outcomes of this relationship are still lacking.

The relevance of better understanding the impact of mitochondrial function in epigenetics cannot be understated given the many ways mitochondrial output has been documented to influence gene expression [9-11]. Novel links between mitochondrial function and epigenetics continue to be unveiled and mechanistic understanding of this relationship is emerging. Tricarboxylic acid (TCA) cycle intermediates such as acetyl-CoA and α-ketoglutarate (α-KG) are substrates or co-factors for enzymes that alter the epigenome, such as the histone acetyltransferases (HATs) and the demethylases [7,12-19]. Thus, mitochondrial dysfunction could, for example, alter the nuclear epigenome through reduced TCA flux. In fact, we first reported that progressive loss of mitochondrial DNA (mtDNA) and the associated changes in TCA output, by ectopically expressing a dominant-negative mitochondrial DNA polymerase (DN-POLG), led to histone hypoacetylation in the nucleus [6]. Using this same cell system, we also demonstrated a direct link between loss of mtDNA and DNA hypermethylation, which we showed was driven by modulation of methionine salvage and polyamine synthesis, both sensitive to changes in TCA cycle flux. We showed that DNA methylation changes occurred predominantly at the promoters of genes that responded to mitochondrial dysfunction, increased progressively over the course of mtDNA depletion, and could be reversed by maintaining NADH oxidation in the mitochondria, even in the context of complete mtDNA loss [8].

While our initial work using the DN-POLG system revealed hypoacetylation of histones in the nucleus as a function of progressive mtDNA loss [6], mechanistic details associated with these effects were not interrogated. Importantly, it remains unknown whether those histone changes are sufficient to alter gene expression and impact functional outcomes. In this work, we used the DN-POLG cells together with a model of chronic mtDNA depletion to establish cause-effect relationships. Using several biochemical, transcriptomics, epigenomics, genetics and pharmacological approaches, we found that histone acetylation loss or gain occurred predominantly on the promoters of differentially expressed genes (DEGs), that even chronic transcriptomic changes were amenable to inducible epigenetic manipulation by supplementation with a TCA intermediate, and that altered histone acetylation status largely preceded gene expression remodeling.

## Results

### Loss ofH3K9ac marks by progressive mtDNA depletion occurs early in the course of mtDNA loss and predominantly in the promoters of differentially expressed genes

Using Western blots and quantitative mass spectrometry, we previously determined that progressive mtDNA depletion in the DN-POLG cells led to histone acetylation changes at specific lysine residues on H3, H2B and H4; H3 acetylation changes were more frequent and pronounced [6]. In addition, we also unexpectedly found that lysine acetylation increased in some histones [6]. Most recently, we described that loss of mtDNA in this cell model was accompanied by progressive transcriptional remodeling [8], providing an excellent platform to interrogate the extent to which the histone acetylation changes were involved in regulating the expression of those genes. To address this question, we performed ChIP-seq in the DN-POLG cells at days 0, 3, 6 and 9. We studied H3K9ac enrichments for several reasons, including the fact that this is primarily a promoter mark and that it was decreased about ~50% at day 9 in the DN-POLG cells [6].

We started by examining the relative *de novo* H3K9ac peak enrichment around the transcriptional start site (TSS) of genes at days 0, 3, 6 or 9 as recommended elsewhere [20]. For quantitative comparisons, tag densities of H3K9ac peaks detected at each timepoint and in each independent biological replicate were normalized against tag densities in matching input DNA control libraries sequenced in parallel (Fig. 1A black lines; see Materials and Methods for details). In all cases, we confirmed consistency across individual sample libraries for genome-wide read count distributions before further study [20]. Libraries included in the analysis were also scored for compliance with proposed quality metrics per the ENCODE Project; these were implemented via the ChiLin quality control pipeline [21], and included fraction of reads in peaks (FRiP), crosscorrelation profiles (CCPs) between replicates for signal to noise ratio measurements and peak coincidence cross-replicates among others. Reproducible peaks used in the analysis were extracted via bivariate ranking of narrow-peak enrichment significance with a 1% Irreproducible Discovery Rate (IDR) cutoff [22]. Using this approach, we found progressive loss of average genome-wide H3K9ac peak densities over time. Notably, the changes in H3K9ac enrichment were significant already at day 3 (Fig. 1A, blue lines), which was unexpected given that neither Western blots or mass spectrometry had shown significant changes at this time [6]. However, these data underscore the sensitivity of ChIP-seq compared to these other approaches for analysis of histone modification abundance. Based on the total number of reproducible peaks and visual inspection of the genomic tracks at each time point, it became clear that peaks were either lost or significantly decreased in the DN-POLG samples over time (Fig. S1A).

**Figure 1.**
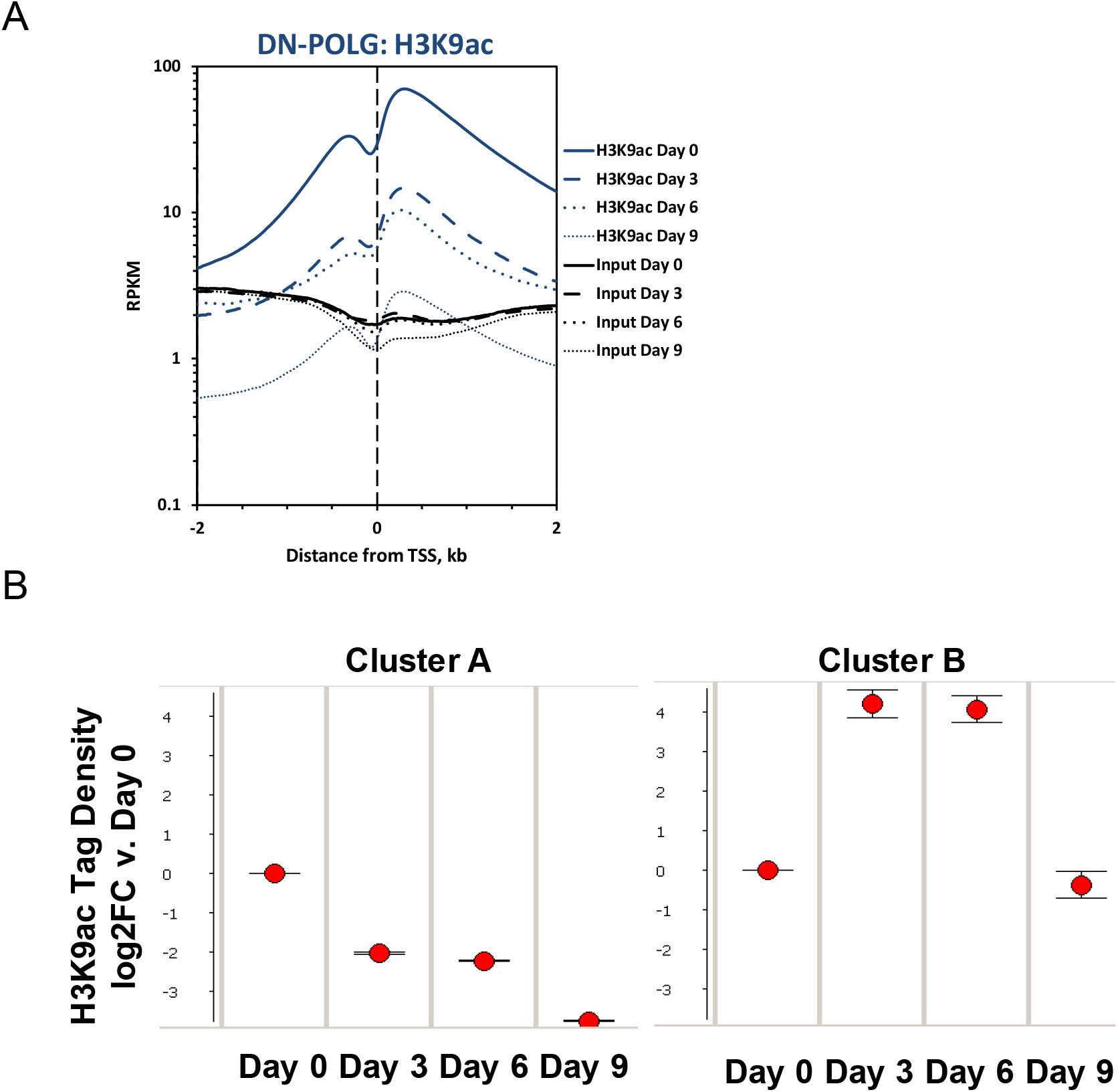
Changes in histone acetylation caused by progressive mtDNA depletion occur in specific loci and are robust already at day 3. (A) Average genome-wide enrichment level of H3K9ac peaks DNA centered around the transcription start site (TSS) of genes. N=2 per time point. Blue line shows data for H3K9ac and black lines for input DNA. (B) Time-dependent patterns of average significantly different enriched H3K9ac peaks relative to day 9 in the DN-POLG cells following hierarchical clustering. Error bars represent standard error of the mean.

We next identified the peaks that showed statistical differences at days 3, 6 or 9 relative to day 0, and performed hierarchical clustering to define how they behaved over time. We found that most peaks decreased as mtDNA was depleted (Fig. 1B, cluster A) while a few increased between days 3 and 6 with levels at day 9 resembling those of day 0 (Fig. 1B, cluster B). Detailed analysis of the peaks following the behavior of cluster B indicated that, in all cases, the peak at the canonical annotated TSS decreased. However, a new peak close to the TSS, but in a non-annotated region of the gene, emerged and was the one increased (see example on Fig. S1B); the relevance and origin of these novel peaks remain unclear. We then cross-referenced the coordinates of the significantly changed peaks with those of the promoters of the 2,854 DEGs identified on the DN-POLG based on RNA-seq [8]. We found that ~73% of DEGs in the DN-POLG cells harbored a change in their promoter H3K9ac peak levels (Table S1). The promoters of 1,995 followed the pattern shown in cluster A, whereas 65 DEGs followed the pattern of cluster B (Table S1). Although we found that modulation in histone peak intensities were detected in ~10,000 genes (Fig. S1C, and Table S2), including those transcribed but not differentially expressed, statistical analysis revealed that the odds ratio of a gene being differentially expressed and having a change in its H3K9ac level was OR=2.41; p<5×10^−90^ (Fig. S1C). These data indicate that modulation of H3K9ac was more prevalent in the promoters of DEGs than of other transcribed genes. The pathways enriched by genes harboring altered H3K9ac levels is shown in Table S1, and revealed that they were involved in key functions affected in our model system. For example, we had previously shown that cholesterol biosynthesis, putrescine metabolism and the TCA cycle, among other pathways, were inhibited by progressive mtDNA depletion [8]. Concomitant to the decreased expression of genes in those pathways was the loss of their H3K9ac promoter mark (Table S1). Taken together, these data suggest that the mitochondrial-driven histone acetylation changes strongly correlate with the differential expression of genes that respond to mtDNA depletion.

### Maintenance of site-specific H3K9ac levels coincide with loss of differential expression of genes and reversal of functional outcomes caused by progressive mtDNA depletion

Resuming NADH oxidation in the mitochondria of the DN-POLG cells through the ectopic expression of NADH dehydrogenase like I (NDI1) and alternative oxidase (AOX) maintained histone acetylation levels even when mtDNA was completely lost [6]. Concomitantly, only 23 genes were shown to be differentially expressed in these cells between days 0 and day 9 by microarrays, while approximately 1,000 genes were identified in the DN-POLG at this same time using this tool [8]. Thus, if histone acetylation changes were required for the differential gene expression, then the promoter H3K9ac levels in the coordinates of the 1,000 genes identified in the DN-POLG at day 9 should be changed. Conversely, H3K9ac enrichment in those same loci should be maintained across time in cells expressing NDI1/AOX. To test this hypothesis, we performed ChIP-seq in the NDI1/AOX-expressing cells following the same procedures employed for the DN-POLG. We found that the average genome-wide H3K9ac peak densities were not changed when cells expressed NDI1/AOX, even when mtDNA was completely lost at day 9 (Fig. 2A). Focusing solely on the promoter coordinates of the ~ 1,000 DEGs, we found that average H3K9ac levels were significantly increased in the NDI1/AOX cells, irrespective of whether peaks followed the pattern of cluster A or B compared to the levels found in the DN-POLG cells (Fig. 2B). Analysis of individual promoters further confirmed that H3K9ac enrichments in the same genomic coordinates were increased in the NDI1/AOX cells compared to the densities identified in the DN-POLG counterparts (representative data on Fig. 2C). It is noteworthy that the average fold-change in the expression of genes in those coordinates in the NDI1/AOX-expressing cells at day 9 were minor and not statistically significant, following the H3K9ac peak densities (Fig. 2D). Thus, we conclude that there is a strong correlation between modulation of the H3K9ac promoter levels and differential gene expression (or lack thereof) in response to mitochondrial dysfunction.

**Figure 2.**
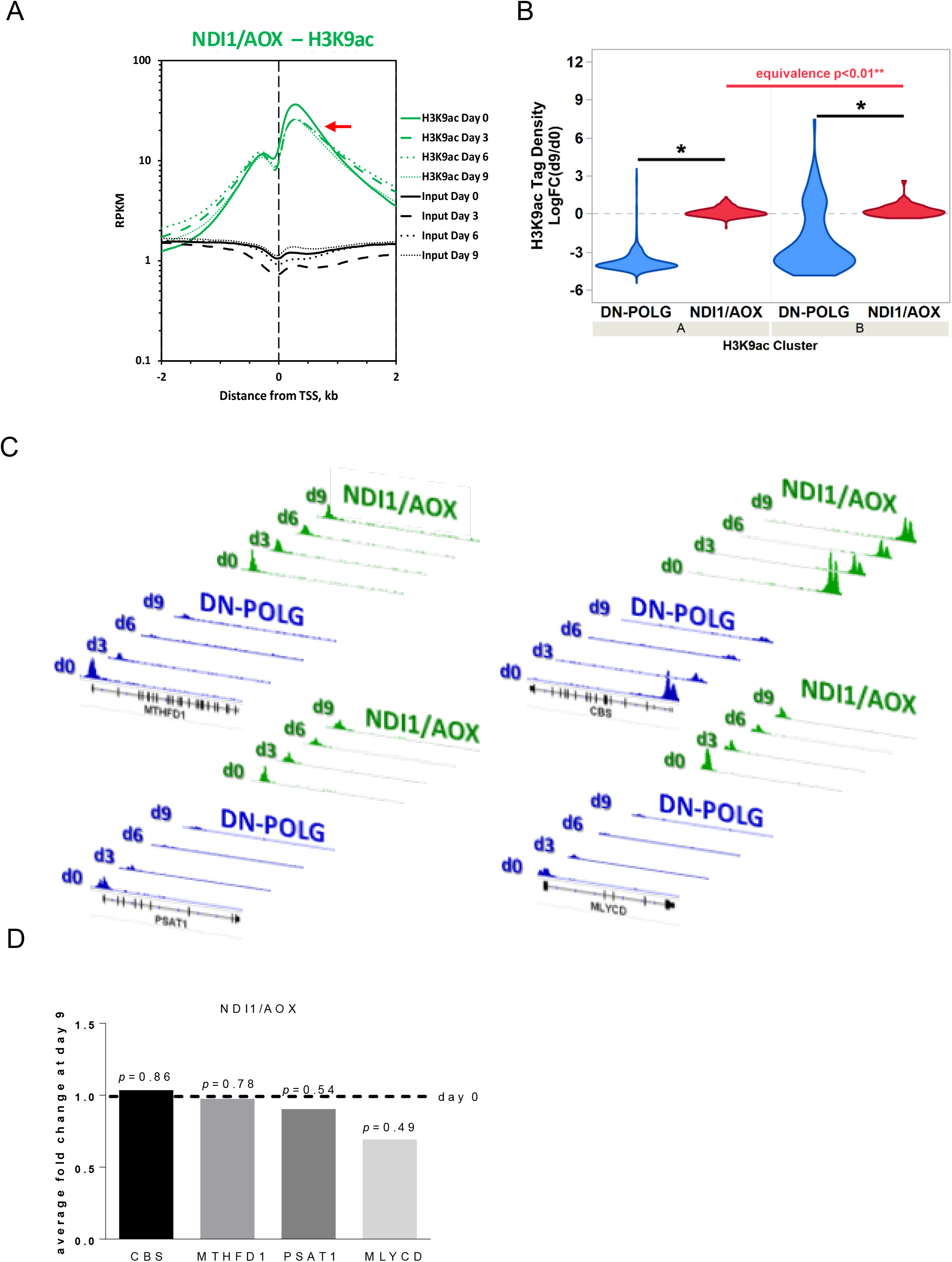
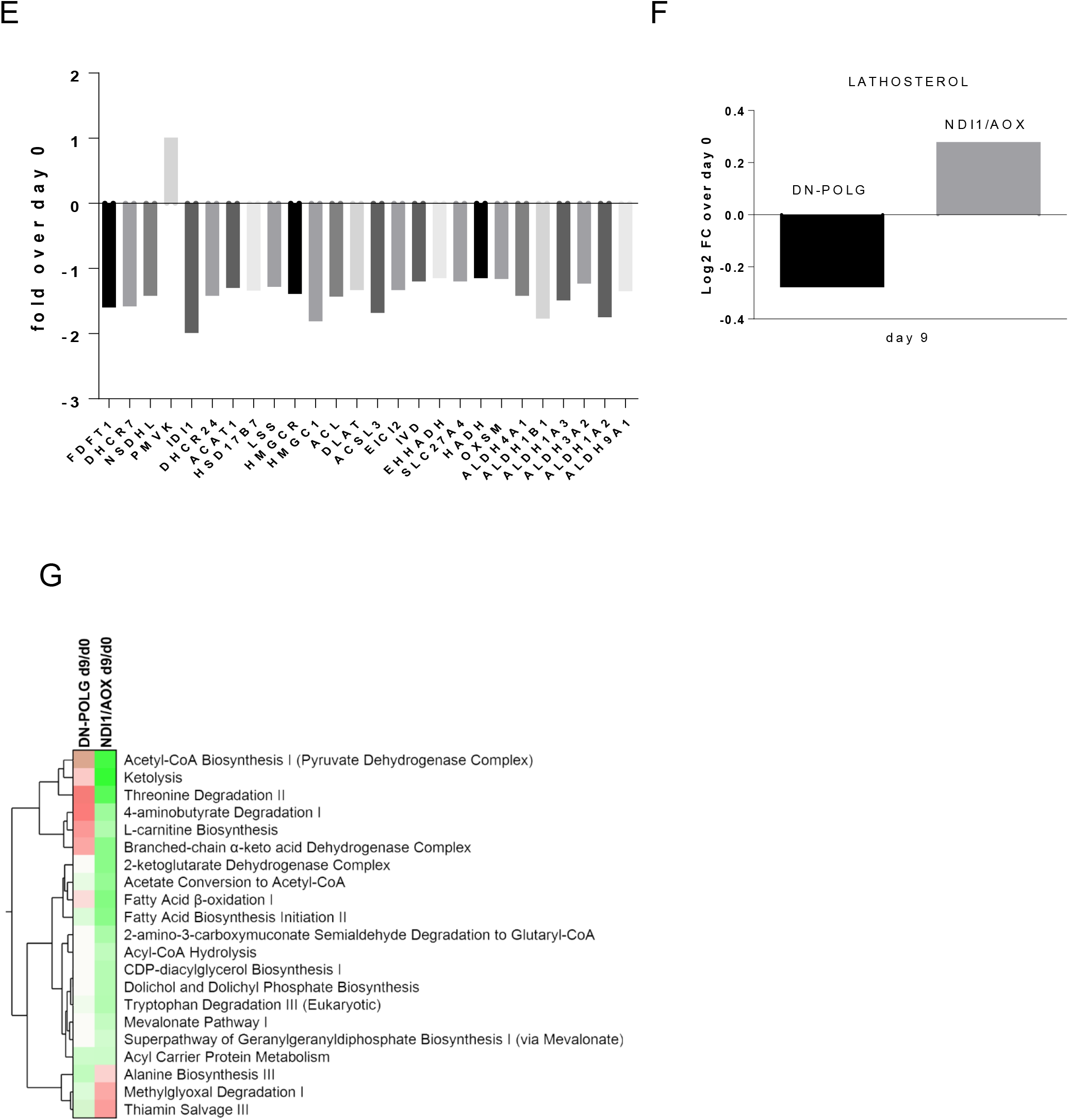
Genetic maintenance of histone acetylation prevents gene expression changes in the promoters of genes responding to acute mtDNA depletion. (A) Average genome-wide enrichment level of H3K9ac peaks (green lines) centered around the transcription start site (TSS) of genes. Black lines show input DNA, which was used for normalization purposes. Red arrow indicates levels of average peak intensity at day 9. N=2 per time point. (B) Log-fold enrichment levels of significantly changed H3K9ac peak tags at day 9 v. day 0 DN-POLG (blue) and NDI1/AOX (red) cells for genes falling on peaks within clusters A or B (as per Fig. 1B). Student’s t-test p<0.01 is depicted by single asterisk (*) between statistical groups; equivalence testing between statistical groups was performed by the two-one-sided tests (TOST) procedure. (C) Graphical representation of the H3K9ac peak densities at the TSS of 4 distinct DEGs identified in the DN-POLG cells (blues); the peaks corresponding to the same genomic loci in the NDI1/AOX counterparts are depicted in green. MTHFD1 (methylenetetrahydrofolate dehydrogenase 1), PSAT1 (phosphoserine aminotransferase 1), CBS (cystathione beta synthase) and MLYCD (malonyl-CoA decarboxylase). Black bar indicates the gene and the vertical bars within depict the exons. (D) Genes within the genomic coordinates depicted in (C) were differentially expressed in the DN-POLG as previously reported [8]; their expression levels as gauged by average fold-changes through microarrays in the NDI1/AOX-expressing cells at day 9 relative to day 0 is depicted. (E) Average fold-changes at day 9 relative to day 0 of genes associated with cholesterol biosynthesis (as per IPA analysis, Table S1) were calculated based on RNA-seq experiments performed in the DN-POLG cells [8]. N=3, statistical significance was gauged adjusting for multiple comparisons (FDR=0.05). (F) Levels of lathosterol identified based on reanalysis of metabolomics data of DN-POLG and NDI1/AOX cells [8]. Log2-fold change was calculated based on N=4 for each cell line comparing levels at day 9 to their respective day 0 counterparts. (G) Heatmap of significantly represented canonical metabolic pathways per Ingenuity Pathway Analysis based on differentially enriched metabolites in DN-POLG and NDI1/AOX cells at day 9 of dox-inducible mtDNA depletion. Color intensity of heatmap blocks corresponds to strength of enrichment [-log(p)>1.3]; red and green hues are representative of pathway enrichment based on predominance of upregulated or downregulated metabolites in the pathway, respectively.

The existence of metabolomics data for the DN-POLG and NDI1/AOX cells [8] provided us with an unique opportunity to determine whether the gene expression changes associated with H3K9ac peak densities had functional outcomes. Many TCA cycle genes had decreased H3K9ac levels, showed inhibited gene expression and decreased metabolite levels in the DN-POLG cells, all of which were rescued in the NDI1/AOX-expressing cells (Table S1 and [8]). We found that, likewise, genes associated with pathways not directly impacted by resumption of mitochondrial NADH oxidation, such as cholesterol biosynthesis, also showed decreased H3K9ac levels (Table S1). All of the genes associated with cholesterol biosynthesis captured in our RNA-seq analyses were downregulated in the DN-POLG cells (Fig. 2E). Only one metabolite directly reflecting cholesterol biosynthesis, lathosterol, was found in our metabolomics analysis [8], but its levels were decreased in the DN-POLG cells and were completely rescued in the NDI1/AOX cells (Fig. 2F). Similarly, other metabolites associated with pathways that could impact cholesterol synthesis were completely rescued in the NDI1/AOX-expressing cells (Fig. 2G). Collectively, these results support the conclusion that the effects of mtDNA depletion on histone acetylation led to functional outcomes, including both at the levels of gene expression and with respect to the metabolic products of the genes/pathways affected.

### Histones are hypoacetylated under chronic mitochondrial dysfunction and are associated with decreased histone acetyltransferase activity

While the genetic manipulation performed in the DN-POLG and NDI1/AOX cells provided unequivocal inferences about the role of mitochondria in epigenetically-driven gene expression regulation, a drawback of these models is that the histone and transcriptome rescues were performed in isogenic but independent cell lines. Ideally, the rescue of the histone mark and downstream effects should be demonstrated in the same cellular background. Hence, we next performed a series of experiments in an unrelated cell line (143B) whose mtDNA had been depleted by exposure to low doses of ethidium bromide (EtBr) [23]; herein these cells are referred to as rho0 and the mtDNA-repleted control counterpart as rho+. Experiments included biochemical evaluation of parameters associated with mitochondrial function (Fig. S2A-D) and the demonstration that 143B rho0 cells have H3K9ac and H3K27ac marks chronically depleted when compared to rho+ controls (Fig. 3A). Overall, these results recapitulate the findings on the DN-POLG model. Also, they suggest the lack of an active compensatory mechanism to maintain histone acetylation when mtDNA is depleted - whether short- or long-term.

**Figure 3.**
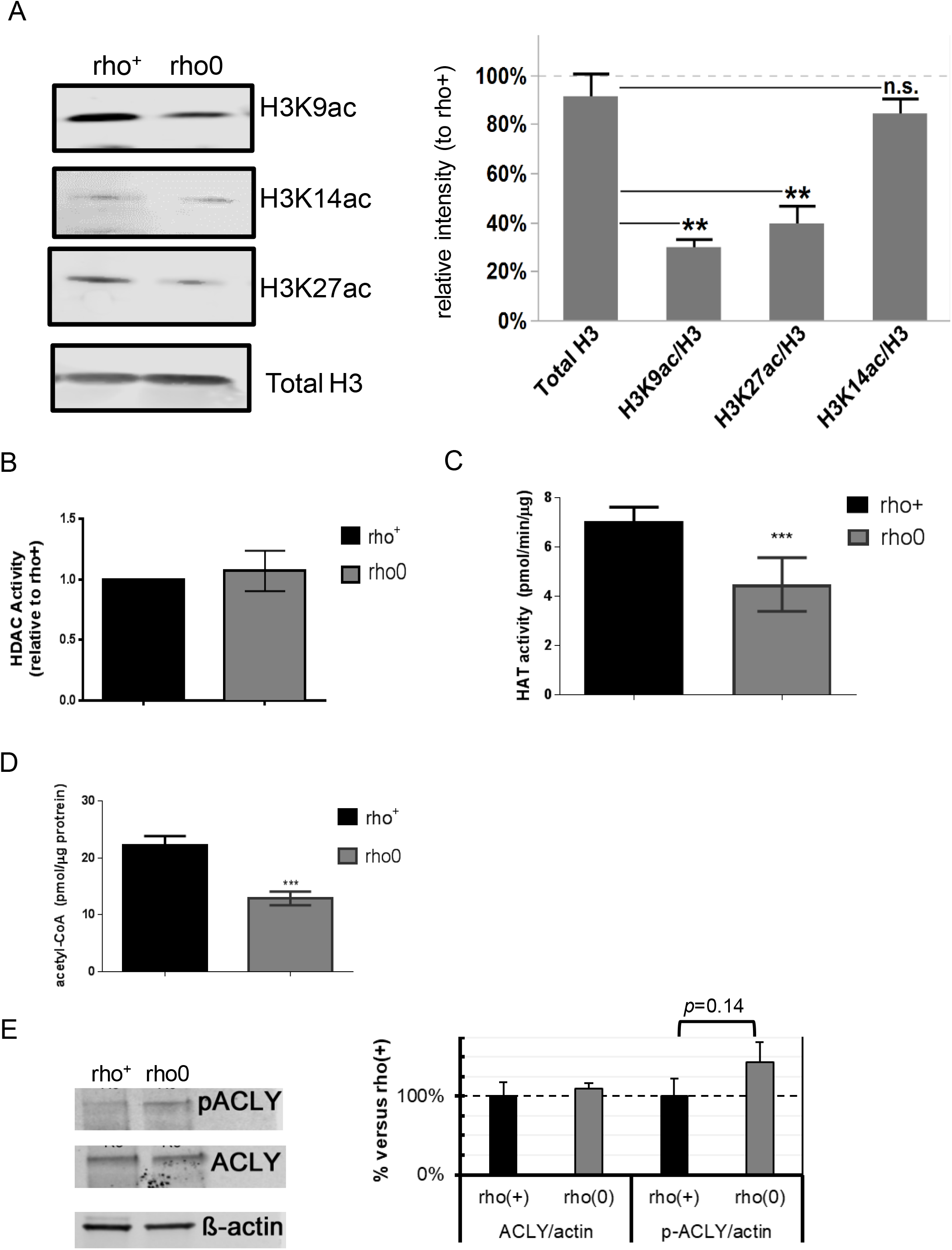
Histones acetylation is decreased in H3K9 and H3K27 in chronically mtDNA-depleted cells due to decreased histone acetyltransferase activity. (A) Representative Western blots of different histone H3 acetylation marks in rho+ and rho0 cells; graph represents average data from N=3 independent experiments. (B) HDAC activity was gauged based on a fluorometric assay using nuclear extracts. N=4. (C) HAT activity was estimated using a fluorescent assay and a standard curve. N=5. (D) Total levels of acetyl-CoA were estimated using deproteinized samples in a fluorescent-based assay and a standard acetyl-CoA curve. N=3 biological replicates. (E) Representative Western blots probing phosphorylated and total ACL levels in 3 independent biological replicates of rho+ and rho0 cells. Graph depicts normalization of the data to actin for each antibody in each of the cell types.

In order to rescue the histone marks in the 143B rho0, it was first necessary to gain insights into the mechanisms associated with their hypoacetylation phenotype. We started out by monitoring global histone deacetylase (HDAC) and acetyl-transferase activities given these are opposing functions that could affect the steady-state level of histone acetylation in cells. While no differences were observed in HDAC activity between rho+ and rho0 cells (Fig. 3B), which ruled out a role for increased deacetylation of histones, HAT activity was ~ 50% reduced in rho0 cells (Fig. 3C). The influence of decreased mtDNA content on HAT enzymatic function was further confirmed in 143B cells freshly depleted of mtDNA by EtBr exposure (Fig. S3A-C) and in mouse embryonic fibroblasts from the TFAM (transcription factor A mitochondria) heterozygote mouse (Fig. S3D), which shows 50% reduction in the amount of mtDNA (Fig. S3E and [24]).

Next, we estimated levels of cellular acetyl-CoA this metabolite is not only the substrate for HAT function and it is primarily generated from mitochondrial-derived citrate by cytosolic ATP citrate lyase (ACL) in mammalian cells. Estimation of total levels of acetyl-CoA using a fluorescent-based assay confirmed significantly decreased levels of acetyl-CoA in rho0 cells compared to rho+ cells (Fig.3D). Given inhibition of ACL was previously shown to decrease acetyl-CoA pools leading to histone hypoacetylation in the nucleus [25], decreased ACL function could mediate the effects of mtDNA depletion on HAT activity and histone acetylation. However, neither ACL protein content or enzymatic activity, as judged by phosphorylation at serine 455 [26], was different between rho+ and rho0 cells (Fig. 3E). Thus, we conclude that depletion of mtDNA negatively influences HAT activity through chronic decreases in total levels of acetyl-CoA but not through impaired ACL function.

Because the 143B rho0 cells were generated by chronic low dose exposure to EtBr, a mutagen, it was formally possible that this treatment mutated HAT genes that impaired their function. Therefore, we compared deep sequenced nuclear DNA from the rho+ or rho0 cells to the reference genome and showed similar changes in both cells (Fig. S4A), ruling out that an increased mutation burden impaired HAT function in rho0 cells. Likewise, no significant decreases in the transcription of HAT or acetyltransferase genes were identified in the rho0 compared to the rho+ by microarrays (Table S3). Protein amounts identified for two main HATs, GCN5 and EP300 (Fig. S4B), which acetylate K9 and 27 residues in histone 3 tails [27,28], were also not different between rho+ and rho0.

If mtDNA depletion influences HAT function by limiting the overall cellular content of acetyl-CoA, as the data above suggests, then modulation of the mitochondrial pool of acetyl-CoA should correspondingly affect HAT activity. Thus, we next set out to test this by exposing rho+ and rho0 cells to different pharmacological agents that modulate mitochondrial acetyl-CoA as previously described [29]. In parallel evaluated HAT activity. We supplemented the medium of rho+ and rho0 cells with dimethyl-α-ketoglutarate (DM-α-KG), a cell permeable form of α-KG that enters the mitochondria feeding the TCA cycle, for 4h as previously described [29]. DM-α-KG increased total cellular acetyl-CoA in rho0 but had no effects on rho+ (Fig. 4A), and restored HAT activity in rho0 cells to levels similar to those in the rho+ counterparts (Fig. 4B). Similarly, exposure of cells to dichloroacetate (DCA), an inhibitor of the mitochondrial pyruvate dehydrogenase kinases (PDKs), also restored HAT activity in rho0 but had no effects on rho+ (Fig. 4C). Conversely, diminishing the mitochondrial acetyl-CoA pool in rho+ cells by exposure to 1,2,3-benzenetricarboxylate (BTC) or perhexiline (PHX), inhibitors of the mitochondrial citrate carrier and carnitine transporter, respectively [29], reduced HAT activity (Fig. 4D). BTC decreased total levels of acetyl-CoA in rho+ (Fig. S5A). Interestingly, pharmacological inhibition of ACL with hydroxycitrate (HC), while diminishing acetyl-CoA and HAT activity in rho+ as also shown by others, had no effects in rho0 (Fig. 4E). These data confirm that the amount of cellular acetyl-CoA, whether decreased in the mitochondria or cytoplasm, influences HAT activity. They also support the notion that the levels of acetyl-CoA in rho0 are already limiting for ACL function.

**Figure 4.**
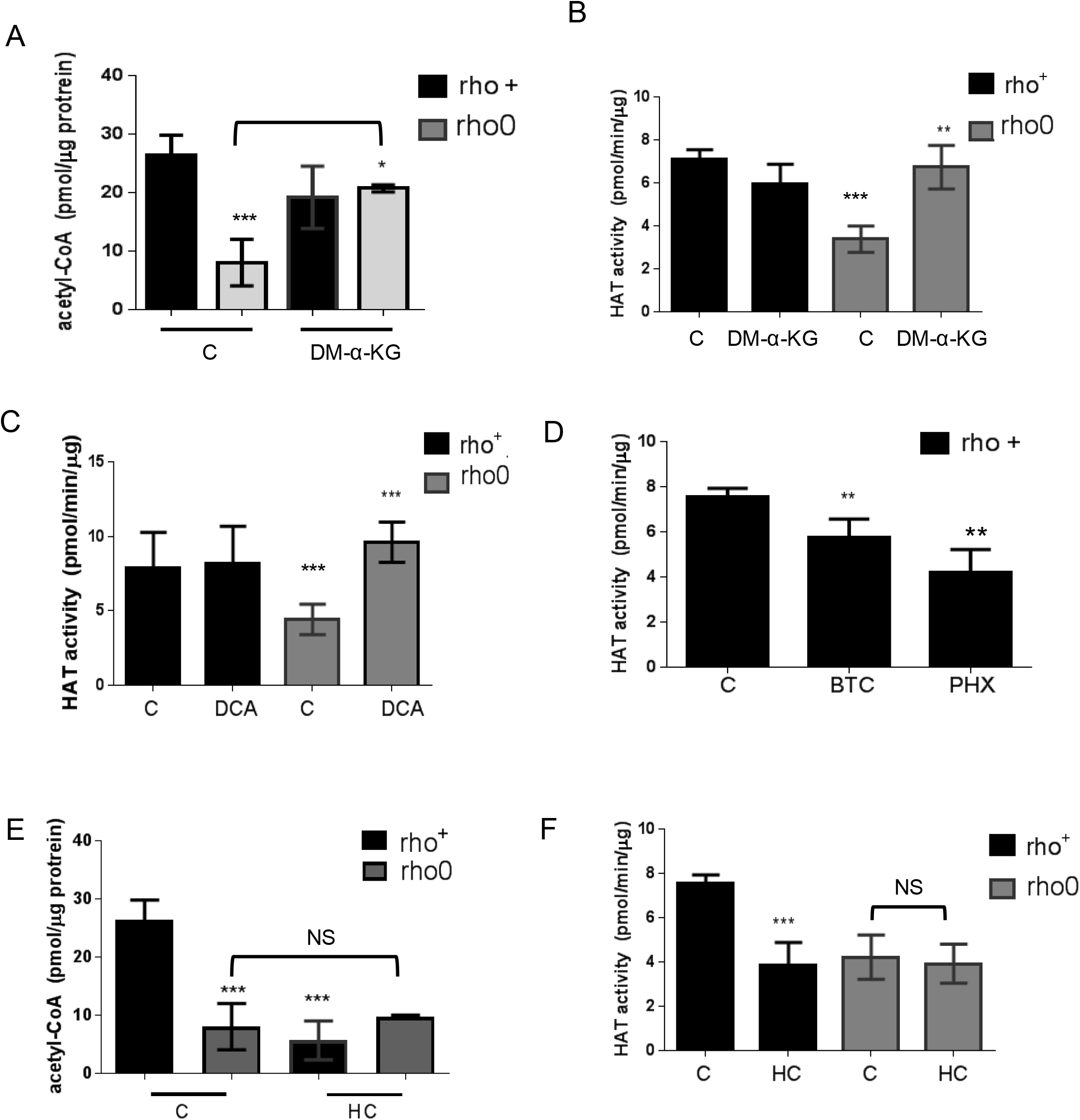
Histone acetyl transferase activity can be modulated by altering the mitochondrial pool of acetyl-CoA. (A) Total levels of acetyl-CoA were estimated prior to or after exposure of rho+ or rho0 cells to dimethyl α-ketoglutarate (DM-α-KG) using deproteinized samples in a fluorescent-based assay and a standard acetyl-CoA curve. N=3 biological replicates. (B) HAT activity was estimated in samples from (A) using an *in vitro* assay. (C) Rho+ and rho0 cells were exposed to the mitochondrial pyruvate kinase inhibitor dichloroacetate (DCA) for 4h prior to HAT activity estimation using a fluorescent-based assay; N=3. (D) Rho+ cells were treated with the mitochondrial citrate carrier inhibitor 1,2,3-benzenetricarboxylate (BTC) or an inhibitor of the mitochondrial carnitine transporter, perhexiline (PHX), and HAT assays performed as in (C). (D) Rho+ or rho0 cells were exposed to hydroxycitrate (HC), an inhibitor of the cytosolic ATP citrate lyase (ACL), for 4h. Total acetyl-CoA levels were then estimated using deproteinized samples in a fluorescent-based assay and a standard acetyl-CoA curve. N=3 biological replicates. (E) HAT activity was gauged in the samples from (D).

### Reversal in locus-specific histone acetylation marks occurs in the promoters of genes whose expression is affected by DM-α-KG supplementation

In addition to modulating HAT activity, the above pharmacological interventions correspondingly altered histone acetylation (Fig. S5B and C) providing the framework to interrogate cause-effects relationships between histone hypoacetylation caused by mtDNA depletion and gene expression regulation in the same cellular context. To address this question, we started by determining the genes differentially expressed between rho0 and rho+ cells by microarrays, which revealed about 3,300 DEGs (Table S3); validation of randomly genes by quantitative real time PCR (qRT-PCR) is shown in Fig. S6A. Then, we supplemented the rho0 cells with DM-α-KG to perform microarrays; DM-α-KG was chosen because it needs to be metabolized in the mitochondria to generate acetyl-CoA [29]. Also, under our experimental conditions it increased acetyl-CoA and HAT activity in rho0 cells but had no effects on rho+ cells (Figs. 4A and B). We found that 596 out of 3,295 DEGs in rho0 cells had their expression changed by the DM-α-KG treatment (Table S4). DM-α-KG partially or fully rescued the directionality of the change for ~70% of the 596 affected genes (Fig. 5A and Table S4), which was noteworthy given the chronic state of transcriptome remodeling in the rho0 cells.

**Figure 5.**
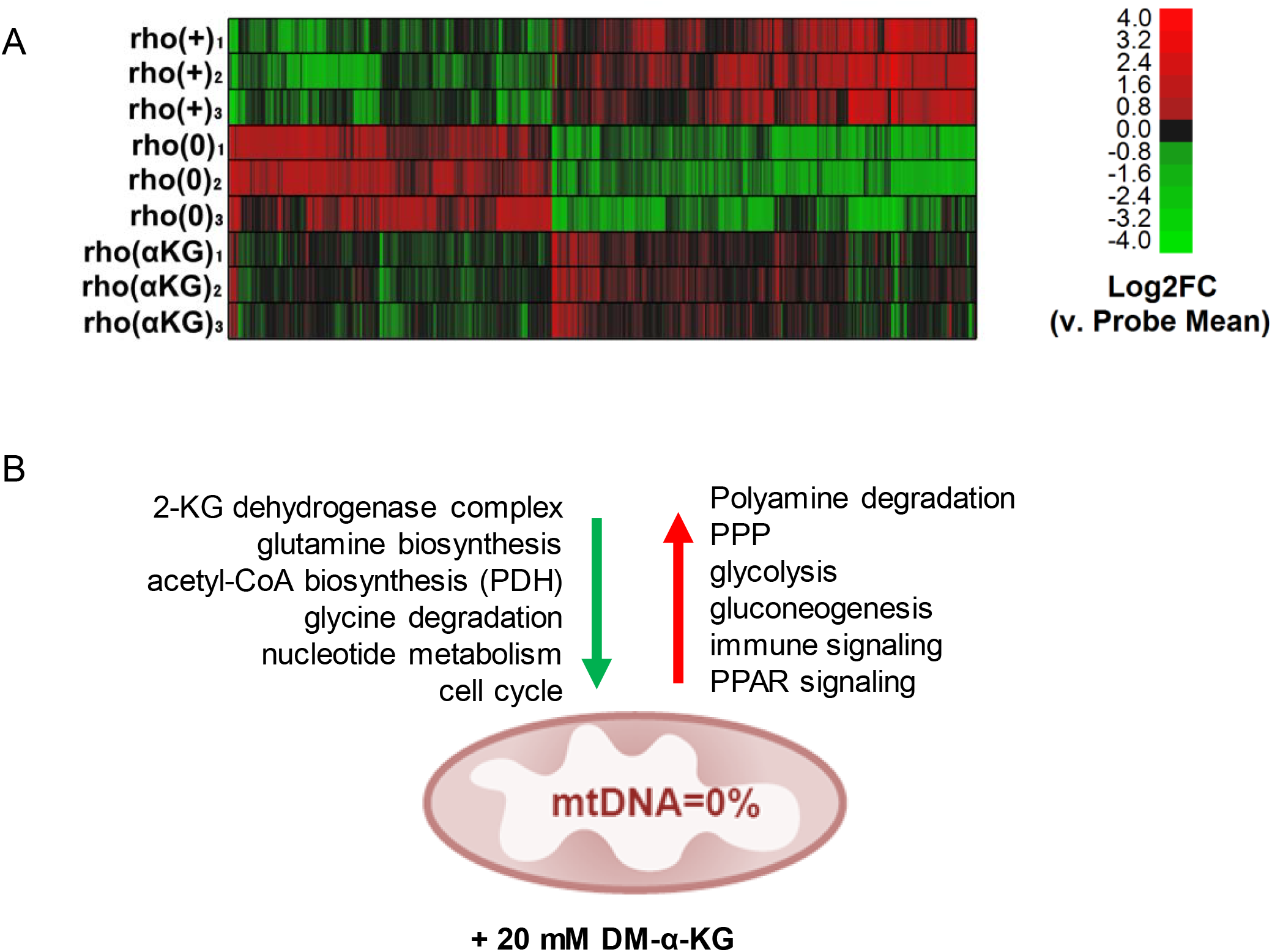
Treatment with DM-α-KG rescues gene expression in the context of chronic mtDNA depletion. (A) Microarrays were performed using RNA from rho+, rho0 or rho0 pre-treated with DM-α-KG; N=3 per sample. Data are presentative of the 596 genes that were impacted by the treatment. (B) Schematic representation of the main transcriptional responses reversed in the rho0 cells by DM-α-KG exposure; green and red arrows indicate downregulated and upregulated signaling pathways, respectively, after treatment.

Pathway enrichment using Ingenuity Pathway Analysis (IPA) revealed that the 596 DM-α-KG-sensitive genes in rho0 cells were broadly involved in metabolic or signaling pathways; a schematic representation of a few enriched pathways is shown in Fig. 5B and the full list can be found in Fig. S6B and C. The transcriptional response of some genes involved in cellular metabolism was not surprising given that many can be affected by changes in acetyl-CoA levels. For instance, SAT1 is an enzyme involved in the catabolism of spermidine and spermine, a reaction that consumes cytosolic acetyl-CoA. SAT1 is known to be regulated transcriptionally and to be dependent on the levels of putrescine, the catabolic byproduct of spermidine/spermine [30]. SAT1 was downregulated in rho0 relative to rho+ cells, consistent with decreased acetyl-CoA availability, but less so upon DM-α-KG exposure (Table S4). Conversely, the sensitivity of genes associated with the immune response and inflammation, which were mostly upregulated in the rho0 cells after DM-α-KG, was unexpected given that there is no reported direct link between these pathways and acetyl-CoA levels or DM-α-KG metabolism. However, connections between these phenotypes and mitochondrial dysfunction do exist.

Interestingly, prediction of upstream regulators of these 596 DM-α-KG-sensitive genes using ChIP-seq data from the ENCODE consortium through Enrichr [31,32] identified the histone acetyltransferase EP300 as the top hit (Fig. S6D). This is consistent with the idea that modulation of mitochondrial acetyl-CoA can influence HAT function, and with the hypothesis that histone acetylation changes regulate gene expression in the context of mitochondrial dysfunction. To directly test whether changes in promoter histone acetylation regulated the expression of those 596 genes, we performed ChIP-seq in rho0 cells prior to and after supplementation with DM-α-KG. We used antibodies against H3K9ac and H3K27ac since both of these marks are associated with promoter regions; H3K27ac marks also maps to enhancer regions [33]. For this analysis, we followed the same rigorous protocol as used for the DN-POLG, first determining the number of peaks identified relative to input DNA. We found a total of 25,340 H3K9ac and 40,885 H3K27ac peaks in rho0 cells and slightly lower numbers after treatment with DM-α-KG: 24,117 for H3K9ac and 30,548 for H3K27ac (Fig. 6A). No effects of DM-α-KG over input DNA were identified (Fig. S7A), which rules out the possibility that the decrease in peak numbers simply reflected changes in normalization parameters. While the changes in the H3K9ac peak numbers were most prominent in gene bodies, those for H3K27ac were also found in intergenic regions. The number of peaks in promoter regions was similar for both marks, irrespective of treatment with DM-α-KG (Fig. 6A).

**Figure 6.**
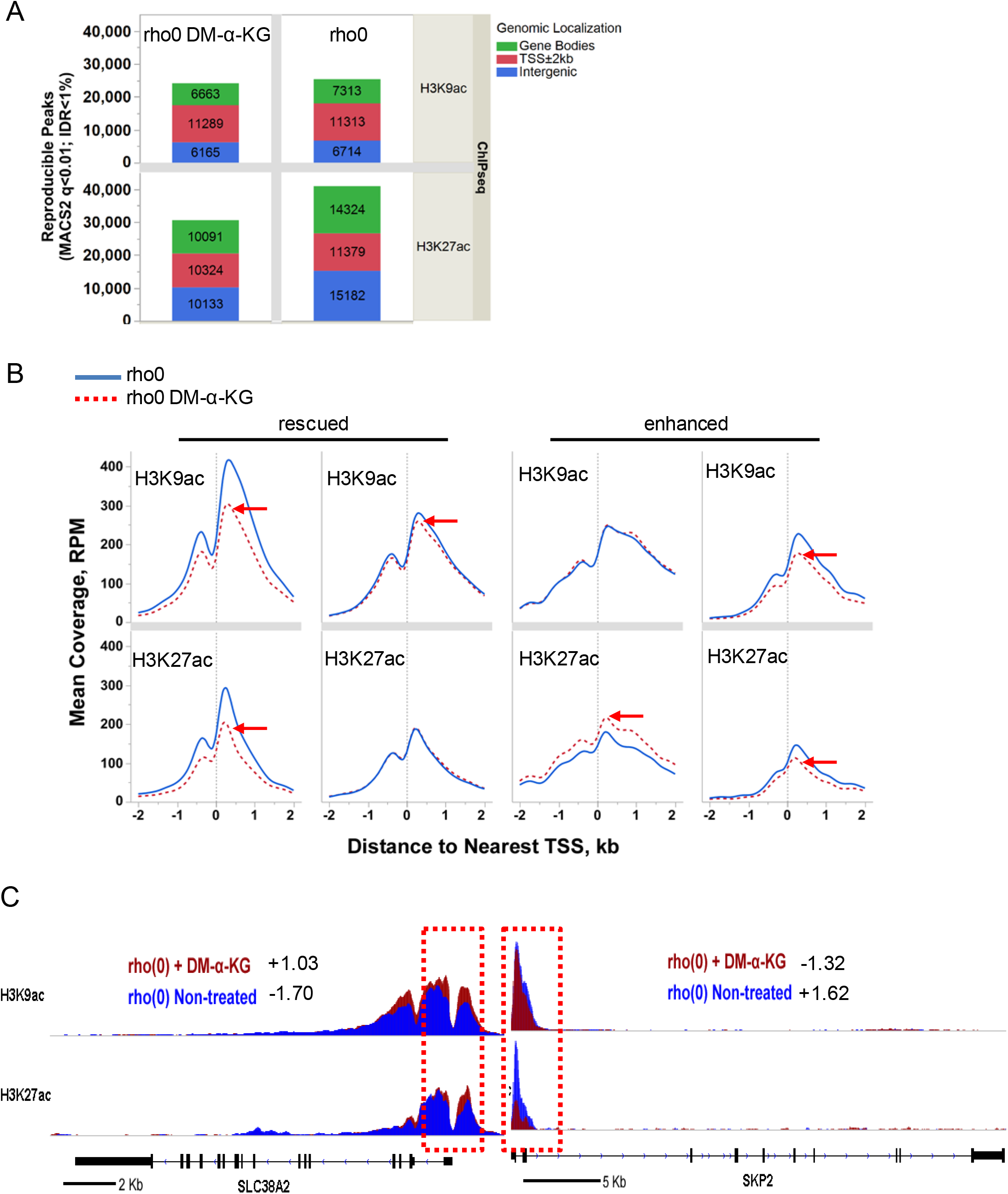
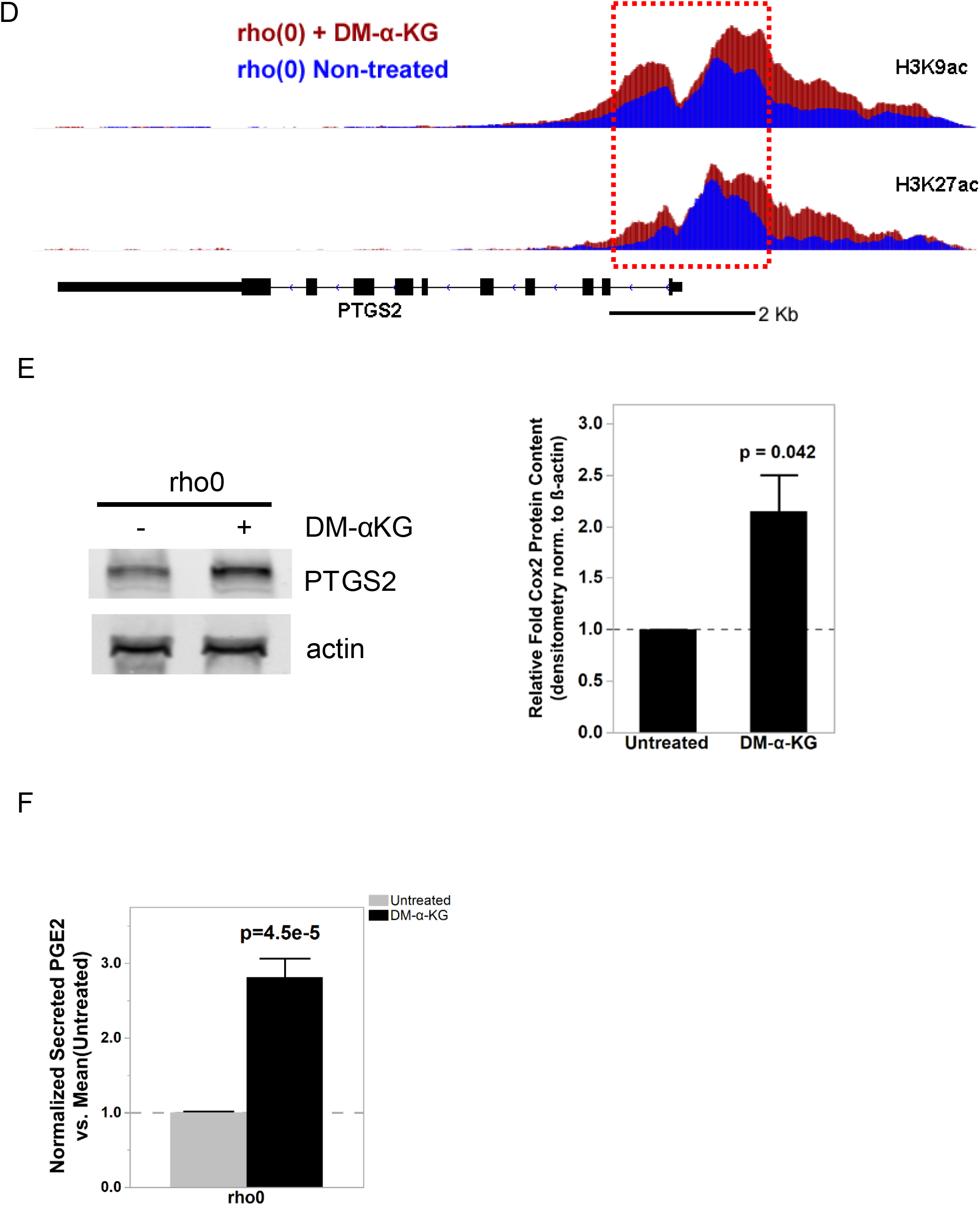
Exposure to DM-α-KG reverses locus-specific H3Kac marks in 143B rho0 cells on the genes affected by the same treatment. (A) Stacked bar plots of genomic localization for H3K9ac (top) and H3K27ac (top) detected reproducible peaks (q<0.01, IDR<1%) by ChIP-seq with respect to gene coordinates in 143B rho0 cells with or without 4h supplementation with 20 mM DM-α-KG in culture. (B) Average enrichment level of H3K9ac and H3K27ac peaks over input DNA centered around the transcription start site (TSS) of 143B rho0 DEGs sensitive to 4h supplementation with 20 mM DM-α-KG in culture. (C) Graphical representation of the H3K9ac and H3K27ac RPM values in two genes differentially expressed in rho0 cells (relative to rho+) with (red) and without (blue) DM-α-KG supplementation; black bar indicates the gene and the vertical bars within depict the exons. Numbers reflect the fold-change relative to rho+ in non-treated or DM-α-KG-exposed rho0 cells. Tracks for SLC38A2 (solute carrier family 38 member 2) and SKP2 (S-phase kinase-associated protein 2) are representative. (D) Same as (C) but for the PTGS2 gene. (E) Representative Western blots of PTGS2 in rho0 cells prior to and after 4h exposure to DM-α-KG; graph shows mean of 3 independent biological replicates. (F) Levels of PGE2 were estimated using ELISA in the supernatant of cells used for (E); N=3. Data were normalized to total protein content.

We started by evaluating H3K9ac or H3K27ac peak status prior to and after DM-α-KG exposure in all 596 genes, independent of whether they were up or down-regulated (Table S4). The behavior of H3K9ac and H3K27ac peaks was similar when evaluating average enrichment across the promoter coordinates of the 596 genes (Fig. 6B and Sig. S7A), although enrichment of H3K27ac was more predictive of changes in gene expression (Fig. S7A). The mean enrichment levels of promoter peaks for the subset of 291 genes upregulated in rho0 cells, but rescued by DM-α-KG, were significantly decreased on their TSSs; similar effects were observed for the 101 genes that were further downregulated in rho0 cells after the treatment with DM-α-KG (Fig. 6B, left panels). These results were consistent with the fact that histone hypoacetylation is associated with decreased chromatin accessibility to transcription factors [34]. Conversely, H3K27ac peak enrichment increased for the 86 genes upregulated in rho0 cells (Fig. 6B, lower right panels). In addition to this average peak intensity, we further analyzed H3K9ac and H3K27ac peaks on specific genes, with representative examples depicted in Fig. 6C. Taken together, these data show that the 596 genes whose expression in rho0 cells that were sensitive to DM-α-KG showed concomitant reversal of their promoter H3K9ac and/or H3K27ac status. These results, in combination with the data on the DN-POLG cells, strongly support a model in which the histone acetylation changes driven by mitochondrial dysfunction regulate gene expression.

We next asked if the locus-specific H3Kac and gene expression changes driven by DM-α-KG supplementation would lead to measurable functional outcomes. To exemplify this relationship, we chose prostaglandin G/H synthase 2, also known as cyclooxygenase-2 (PTGS2 or Cox2), as this gene was decreased in rho0 versus rho+ cells, although its expression was completely recovered in rho0 relative to rho+ cells after DM-α-KG treatment (Table S4). PTGS2 converts arachidonate to prostaglandin E2 (PGE2), which can be measured in the tissue culture supernatant. Concomitant to the increased gene expression in rho0 cells after DM-α-KG treatment was the significant increase in promoter abundance of the H3K9ac and H3K27ac marks (Fig. 6D). Changes in PTGS2 mRNA in rho0 cells led to a parallel increase in protein (Fig. 6E and Fig. S7B), which resulted in significant increases in the levels of secreted PGE2 after exposure to DM-α-KG (Fig. 6F). No significant changes in protein or PGE2 levels were observed in rho+ exposed to DM-α-KG (Fig. S7C and D). Therefore, these data strongly link mitochondria-driven changes in histone acetylation and gene expression to functional outcomes.

From an epigenetic perspective, DM-α-KG could affect the methylation status of the epigenome since α-KG is a co-factor of enzymes that drive demethylation reactions, e.g. the Ten Eleven Translocation (TET) enzymes [35]. Interestingly, we recently showed that the DNA of these rho0 cells is primarily hypermethylated, affecting 621 DEGs compared to the rho+ cells [8]. Even though we showed that DNA methyltransferase activity was increased in the rho0 cells [8], it was formally possible that DM-α-KG treatment could decrease DNA methylation by promoting TET activity in turn affecting gene expression in our experiments. To gain insights into this possibility, we cross-referenced the coordinates of the 596 genes that were affected by DM-α-KG with the 621 differentially methylated genes in rho0 cells. We reasoned that if DM-α-KG affected DNA demethylation in a way that would reverse gene expression, then the loci associated with those genes should start out by being hypermethylated in rho0 cells. Out of the 596 DM-α-KG-sensitive genes, 101 were differentially methylated in rho0 relative to rho+ cells (Table S5), but only 40 of these started out as being hypermethylated. Thus, if differential DNA methylation also played a role in effects associated with DM-α-KG treatment, only a small portion of genes that reversed expression (~10%) may have had their transcription influenced by this epigenetic modification.

## Discussion

Mitochondrial function is key to organismal health, and it is now accepted that mitochondria affect cellular physiology through mechanisms beyond bioenergetics and ROS. However, the means through which modulation of mitochondrial metabolism can effectively change biological outcomes is still under investigation. Because various mitochondrial TCA metabolites are either substrates or co-factors of enzymes that impact the epigenome, presumably mitochondrial dysfunction or mitochondrial metabolic rewiring can impact epigenetic regulation of gene expression. Despite these assumptions, no report has demonstrated the requirement of mitochondrial function for long-term maintenance of chromatin acetylation. Moreover, to our knowledge there is no previous available evidence showing that changes in histone acetylation can be influenced by the modulation of the mitochondrial output of acetyl-CoA in way that can affect HAT function and the expression of genes within the nucleus.

Here we report that under mtDNA depletion, steady state levels of acetyl-CoA and histone acetylation were decreased, which was observed both when mtDNA was progressively lost (DN-POLG cells) or chronically depleted (143B rho0). We also showed that these effects were associated with the mitochondrial output of acetyl-CoA, which in turn influenced HAT activity in a reversible way, overall contributing to the regulation of gene expression in the nucleus. Several of these findings were unexpected. Firstly, the maintenance of hypoacetylated histones under chronic mitochondrial dysfunction suggest that, unlike the response to loss of ACL function [36], loss of acetyl-CoA associated with mtDNA depletion is not compensated for. At least in terms of gene expression, we identified no obvious means to increase acetyl-CoA production, including through acetate or lipid metabolism or enhanced ALC transcription, (Table S3). Alternatively, some level of compensation may exist, but perhaps maintenance of histone acetylation may be secondary to other non-histone proteins, particularly in the context of mitochondrial dysfunction. This is a compelling possibility considering that acetylation is the second most abundant post-translational modification of proteins in cells, with ~90% of metabolic enzymes estimated to be regulated by it [37], including 20% of the mitochondrial proteome [38]. Along those lines, others have proposed a role for histones as reservoir of acetate that can be used for metabolic processes under specific conditions, depending on the needs of the cell [39]. It is possible that maintenance of cellular or even mitochondrial function itself rely on a steady state level of acetylation reactions above a minimal functional threshold.

Secondly, the findings that HAT activity was influenced by the mitochondrial output of acetyl-CoA has not been previously reported. HATs can be regulated by substrate/co-factor availability, protein-protein interactions and post-translational modifications, including acetylation [40]. While it is known that some HATs have kinetic properties that would allow them to respond to fluctuations in the levels of acetyl-CoA [41,42], our *in vitro* HAT assays were done in the presence of excess acetyl-CoA, making it unlikely that substrate availability was limiting. Nevertheless, in the cellular context this could still be formally possible. It is likely that the decreased levels of acetyl-CoA under our experimental conditions may have changed the post-translational state of some HATs or other proteins with which they interact within the chromatin context. Irrespective of the means, the effects of pharmacological manipulation of the mitochondrial pool of acetyl-CoA both in the rho0 and in the rho+ cells on HAT function unveiled a novel mechanistic link between mitochondrial metabolism and histone acetylation. It is interesting that we found here and in our previous work [6] that changes in abundance of lysine acetylation only occurred in some residues on H3, H2B or H4 when mtDNA was depleted. This is contrary to previous work that showed that glucose deprivation led to global histone deacetylation in H3, H2B and H4 [43]. It is not clear why an overall cellular decrease in acetyl-CoA provided by mitochondrial dysfunction would affect some lysine residues and not all. Interestingly, fluctuations in acetyl-CoA were shown not only to affect global levels of histone acetylation but also the pattern of acetylated residues, by altering lysine acetylation preference by some HATs [44]. It is possible that cellular metabolism is impacted in fundamentally different ways when mitochondria are dysfunctional compared to glucose withdrawal, overall affecting HATs (through post-translational modifications and/or protein-protein interactions) in distinct ways.

Thirdly, the genetic rescue of the TCA cycle in the HEK293 DN-POLG cells, together with our DM-α-KG supplementation experiments in the 143B rho0 cells, provide the most compelling data to connect mitochondrial function with changes in histone acetylation, gene expression and physiological outcomes with cause-effect relationships. This is supported by recent work from another group that showed that metabolism of acetate, pyruvate and fatty acid, specifically through the mitochondria, influence H3K27ac levels and promote cell adhesion [45]. Under our experimental conditions, the fact that expression of 20% of the DEGs was rescued by a short 4-hour exposure to DM-α-KG was remarkable given that these cells were in a state of chronic mitochondrial dysfunction with a well-established pattern of gene expression. The modest changes observed in the levels of histone acetylation after treatment with DM-α-KG may explain why only a fraction of the genes were affected. It is possible that longer treatments with DM-α-KG would have more pronounced effects. Alternatively, perhaps only some genes would be responsive to such intervention. Although our data suggest a role for histone acetylation for the regulation of gene expression, DM-α-KG supplementation or restoration of TCA flux (as in the NDI1/AOX cells) by increasing cellular levels of acetyl-CoA could also impact generalized protein acetylation, including that of transcription factors. Likewise, the overall metabolic rescue provided by these interventions could play a direct role in turning off the signals for differential transcription. Considering that we found that both metabolic and other categories of genes were differentially expressed, we do not favor this possibility.

About 70% of DEGs had an altered H3K9ac mark in the 9-day course of mtDNA depletion in the DN-POLG system, which was remarkably apparent already at day 3. Since the genomic changes at day 3 precede signs of measurable mitochondrial impairments [6], these data suggest that histone acetylation is highly sensitive to changes in mitochondrial metabolism and may prove eventually to be a biomarker of mitochondrial dysfunction. In addition, because broad range transcriptional changes were not identified at day 3, these data suggest that remodeling of the histone acetylation landscape may be required for the full-fledge change to the transcriptional program in response to mtDNA loss. Notably, not all DEGs had changes in their promoter H3K9ac marks. Likewise, many genes that were transcribed but were not differentially expressed did show changes in H3K9ac abundance, which was also the case when we monitored modulation of DNA methylation [8]. Although we found that a gene was statistically more likely to be differentially expressed if it had a change in its measured epigenetic status, what these data highlight is that the epigenetic changes by themselves are not sufficient for the differential expression of genes. Rather it is possible that the effects of mitochondria on the epigenome may serve to poise the genome to respond to additional stimuli. The recruitment of transcription factors, epigenetic writers, readers and erasers may ultimately lead to the transcriptional response.

In summary, our studies uncovered a significant and yet unappreciated means through which mitochondrial function can impact the cell. Additional experiments to try to identify which HATs and dissect the exact means through which they are affected by mitochondrial dysfunction will be fundamental in defining more in depth the mechanistic link between mitochondrial metabolism and histone acetylation. However, our findings provide a new foundation to address phenotypic heterogeneity and tissue-specific pathology associated with mitochondrial dysfunction, including that which is caused by mitochondrial genetic diseases. These results may not only be relevant to the pathophysiology of mtDNA depletion syndromes but also to the effects of environmental agents that lead to mtDNA loss, such as nucleoside reverse transcriptase inhibitors (NRTIs) that are used to treat, for instance, HIV infections. They may also be pertinent to the effects of many antibiotics that, by inhibiting mitochondrial protein synthesis, lead to functional effects comparable to mtDNA loss. Our finding that supplementation with DM-α-KG can rescue histone acetylation and the differential expression of many genes associated with chronic mitochondrial dysfunction could provide the underlying mechanism for the improvement in muscle strength in a mouse model of mitochondrial myopathy by a 5-month supplementation with 5 mM α-KG recently reported [46]. Assuming that such a mechanism is broadly applicable *in vivo*, this could eventually prove to be a novel and intriguing approach for therapeutic purposes. While the concentration of DM-α-KG utilized in our and other studies [46] is well beyond physiological levels, these proof-of-concept experiments open challenges and opportunities regarding metabolic modulation of mitochondrial function to impact health and disease.

## Experimental Procedures

### Cells, cell cultures and experimental conditions

The DN-POLG and NDI1/AOX cells were described recently and maintained according to the previous published work [6]. The osteosarcoma cell line 143B and the rho0 derivative were graciously obtained from Dr. Eric Schon at Columbia University. To generate freshly mtDNA depleted cells, 143B were exposed to EtBr (50 ng/mL) for 2 weeks and subcultured for at least one week without this agent prior to utilization. TFAM MEFs were a kind gift from Dr. Gerald Shadel (Yale School of Medicine). All cells were routinely grown in DMEM high glucose (4.5 g/L) supplemented with 10 mM pyruvate, 50 μg/mL of uridine, 10% FBS and 1% penicillin/streptomycin under 37°C and 5% CO_2_. TFAM MEFs media did not contain uridine. All experiments were done with confluent cell cultures. Acetyl-CoA manipulations using HC, PHX, DCA, and BTC were performed as previously described [29] with DM-αKG concentrations between 5-20 mM; the pH of final drug concentrations was adjusted to 7.0 using NaOH.

### ATP, acetyl-CoA, NAD^+^/NADH ratio and H_2_O_2_ measurements

Cells were pelleted and resuspended in 3.5% of perchloric acid. Lysates were then sonicated on ice for 30s, flash frozen in liquid nitrogen, thawed on ice and then span at 5000xg at 4°C for 10 min. Supernatants were then collected, neutralized to pH ~7.2 with 1M KOH and 50 μL used for the ATP and acetyl-CoA assays (BioVision) following the manufacturer’s instructions. Levels of these metabolites were estimated based on standard curves; data were normalized to protein content obtained from parallel cell cultures. The NAD+/NADH ratio was determined using an equal number of cells (10,000) per cell type with a kit from Promega. Amplex Red (Invitrogen) was used to detect the levels of H_2_O_2_ released in medium as previously described [47].

### Histone Preparations and Western Blots

Histones were purified from nuclear lysates using trichloroacetic acid and acetone as described by others [48]. Relative amounts of modified histones were assayed by SDS-PAGE immunoblotting in multiple independent biological replicates (N≥3 per cell derivative). Total histone protein content per sample was estimated from parallel immunoblots to detect total H3 signal using a pan-specific primary antibody as loading control. All histone antibodies were obtained from Active Motif (Millipore) or LiCor (secondary antibodies). For PTGS2, antibodies were purchased from Santa Cruz Biotechnology; primary antibodies raised against total or phosphorylated ACL protein were obtained from Cell Signaling.

### HAT and HDAC activities

Cells were lysed and the nuclear fraction enriched using differential centrifugation. HAT activity was gauged using 1 μg of nuclear lysates and a fluorometric assay (Biovision) following manufacturer instructions. HDAC activity was assayed using 1 μg of nuclear lysates and a fluorometric kit available from Enzo Life Sciences following manufacturer’s instructions. Data were normalized to protein content.

### PGE2 content

Levels of secreted PGE2 were estimated using an ELISA kit following the manufacturer’s instructions (Cayman Chemical). Data reflect 3 independent biological replicates and were normalized to total protein content.

### ChIP-seq and data processing

Chromatin immunoprecipitations were performed with modifications as recommended elsewhere [49,50]. Briefly, cells grown in adherent monolayers (20-30 million per individual replicate, N=2) for each of the 143B rho+ and rho0 derivatives, or the DN-POLG and NDI1/AOX derivatives, were crosslinked by addition of paraformaldehyde at a 1% (v/v) final concentration directly to culture media followed by 8-min incubation at room temperature; chemical cross-linking was quenched by further supplementation with glycine at 125 mM. Cross-linked cells were scraped off the plates, washed in PBS, pelleted by centrifugation at 4,000×g and 4°C for 10 min, homogenized in 2 mL of lysis buffer containing protease inhibitors (Halt [Thermo-Fisher Scientific]), and sheared with a temperature-controlled BioRuptor instrument with high-power settings at 4°C for 20-25 1-min pulses (50%-duty cycle) of ultrasonic shearing. Sheared cell homogenates from 2 biological replicates containing 1-10 μg DNA each were used to extract control (10% input DNA) and ChIP DNA templates with antibodies against H3K9ac or H3K27ac (ActiveMotif). Each template was ligated and amplified into sequencing libraries using different single-indexed adapters (TruSeq RNA v2, Set A [Illumina]). Each individual library was PCR-amplified in 2 technical replicates for no more than 10 cycles (screened per sample by RT-PCR as the number of cycles to reach the inflection point in log-scale amplification curves); afterwards, duplicate PCR reaction volumes per sample were collected for DNA purification with size-selection by double-sided 0.6X-0.8X SPRI (expected fragment size: 250-450 bp inclusive of sequencing adapters) using AMPureXP magnetic beads (Beckman-Coulter). Upon surveying library quality control, which was performed in 4-plexity runs of 20-30 million 2×35nt paired-end reads in a MiSeq system [Illumina], samples were sequenced in 8-plexity runs using a NextSeq 500 system with high-output flow cells [Illumina] following the manufacturer’s protocols. Adapter 3’ sequences were removed from raw ChIP-seq reads after filter for quality phred scores>20, followed by 5’ trimming of bases 1-10 of each read. The resulting paired-end 2×25nt reads were aligned to the hg19 human reference genome [Genome Reference Consortium GRCh37 from February 2009] [51] with Bowtie2 –sensitive-local settings and 1,000-bp maximum fragment length (–D 15 –R 2 –N 0 –L 20 –i S,1,0.75 –X 1000) [52]. Within-sample consistency between replicative sequencing runs from individual ChIP templates was confirmed based on Pearson pairwise correlation scores of log-transformed uniquely mapped and de-duplicated aggregate RPKM values across ~18,000 non-overlapping 10-Kb genomic regions centered around gene TSS. To assess levels of between-replicate concordance of H3K9ac or H3K27ac peak enrichment in each experimental group, we implemented ChiLin [21] with paired-end mapping (Bowtie 2) [52] and narrow peak-calling modes (MACS2, FDR q<0.01 v. input DNA) [53]. Peak calling was followed by estimation of reproducible peak numbers via bivariate ranking of narrow-peak enrichment significance at the 1% Irreproducible Discovery Rate (IDR) level [22]; when detected across samples, overlapping peaks were merged into consensus genomic tags of differential H39ac or H3K27ac occupancy. Lists containing all consensus genomic loci detected across experimental groups, with differential H3K9ac or H3K27ac peak enrichment, and passing the IDR<1% threshold, were assembled into a conglomerate set of reference genomic tags for further data analysis using SeqMonk, version 37.1 (Andrews, S. SeqMonk, 2007: http://www.bioinformatics.babraham.ac.uk).

### Lysine acetyltransferase gene mutation analysis

To find SNPs in rho0, we used DNA reads sequenced with a MiSeq system [Illumina] and aligned to the hg19 human reference genome [Genome Reference Consortium GRCh37 from February 2009] [51]. The human genome sequences were downloaded from ENSEMBL ftp site [54]. Alignments were performed using Bowtie2 [52]. We then parsed alignment results from the coordinates of the 17 genes from Lysine Acetyltransferase family, each SAM/BAM extension file into pileup data using samtools [55]. The coordinates of each gene on hg19 were taken from NCBI’s Gene database [56]. A SNP was called if over 80% of reads containing a change from original nucleotide aligned at call position. A Perl script was developed to perform the parsing CIGAR field of the mpileup file into Variant Call Format (VCF), to extract all SNPs and INDELs. No INDELs were found. The final comparison of sets of SNP positions between the samples was done using BedTools [57] and Linux terminal commands (bash).

### Gene expression experiments by microarray technology and data analyses

For microarrays analysis of gene expression, the Affymetrix Human Genome U133 Plus 2.0 GeneChip arrays were used. Samples were prepared as per manufacturer’s instructions using total RNA. Arrays were scanned in an Affymetrix Scanner 3000 and data was obtained using the GeneChip Command Console and Expression Console Software (AGCC; Version 3.2 and Expression Console; Version 1.2) using the MAS5 algorithm to generate CHP-extension files. Analysis of variance (ANOVA) was used to identify statistical differences between means of groups at α<0.05 level among HG-U133 Plus 2.0 probe sets unambiguously mapped to UCSC known gene transcripts. Gene expression patterns were determined empirically from log_2_-transformed expression fold-changes by unsupervised hierarchical clustering (Ward’s distance metric) using JMP software (Version 11).

## Acknowledgments

We thank the staff at the Core Facilities at NIEHS and NIH (Epigenetics and Genomics) and the critical reading of the manuscript by Drs. Matthew Longley and Robert Petrovich (NIEHS). This research was supported by the Intramural Research Program of the NIH, National Institute of Environmental Health Sciences.

## Author Contributions

OAL performed and analyzed genomics, prostaglandin and qRT-PCR experiments, DG maintained cell cultures, performed HAT and Western blot assays, MS helped set up HAT assays and performed HDAC analysis, TW helped with bioinformatics, TCW performed metabolite analysis in 143B cells, DGS performed the Seahorse experiments, GR performed the mutation analysis of the lysine acetyltranferase genes, NC was involved in the conceptualization of the HEK293 experiments, RPW participated in writing of the manuscript, JHS conceptualized the experiments, helped interpreted the data (with OAL) and wrote the manuscript.

## Data accessibility

Genomics data for this publication have been deposited in the NCBI’s Gene Expression Omnibus and are accessible through GEO Series accession number GSE100134.

## Hyperlinks to Supplemental Tables

“https://orio.niehs.nih.gov/ucscview/Santos/Table_S1-H3K9ac_DEGs_and_IPA_in_DN-POLG.xlsx”

“https://orio.niehs.nih.gov/ucscview/Santos/Table_S2_H3K9ac_at_9541_Expressed_Genes.xlsx”

“https://orio.niehs.nih.gov/ucscview/Santos/Table_S3-differentially_expressed_genes_in_rho0.xlsx”

“https://orio.niehs.nih.gov/ucscview/Santos/Table_S4-genes_affected_by_aKG_in_rho0.xlsx”

“https://orio.niehs.nih.gov/ucscview/Santos/Table_S5-DMEGs_affected_by_aKG_in_rho0.xlsx”

---

**Fig. S1.**
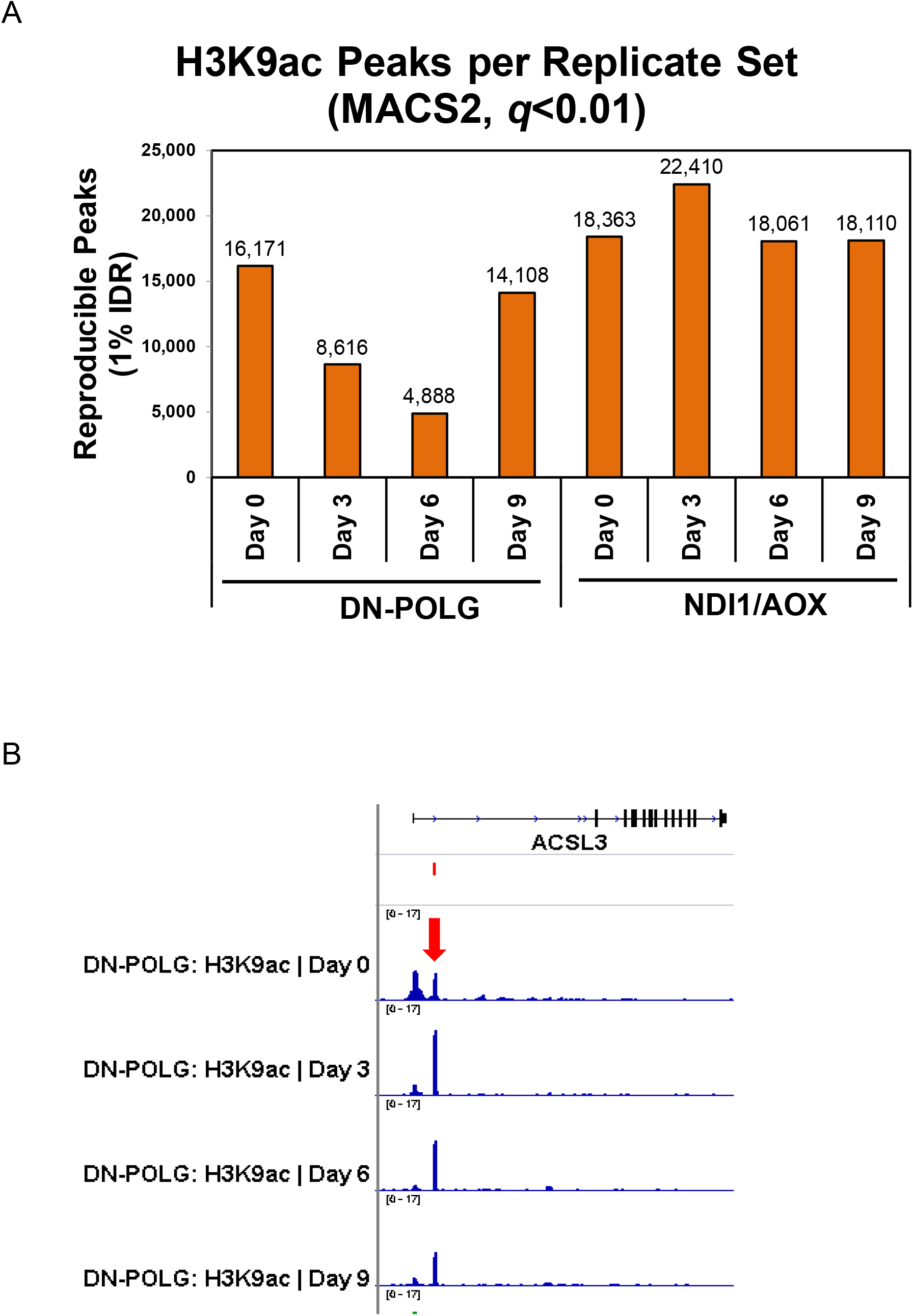
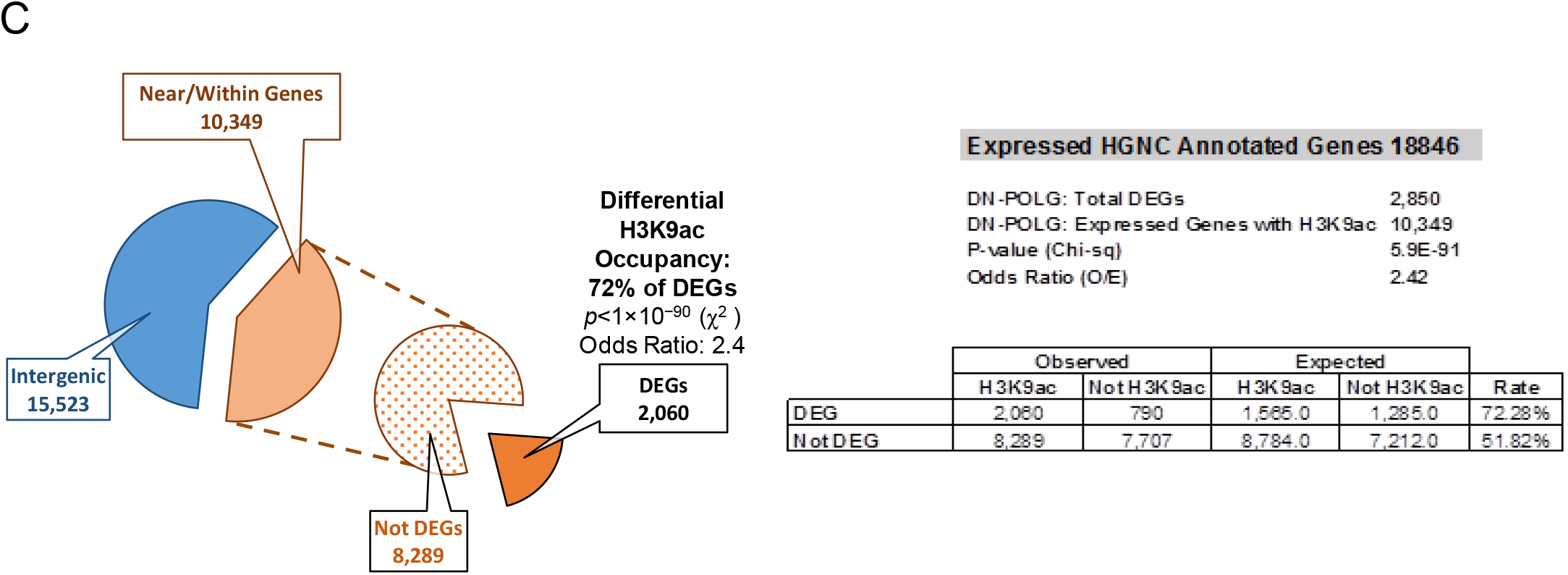
Detection of H3K9ac enrichment peaks during the 9-day timeframe of dox-induced mitDNA depletion by ChIPseq. (A) Number of reproducible H3K9ac enrichment peaks v. input DNA genome-wide at days 3, 6 or 9 v. day 0 (MACS2, q<0.01, 1% IDR); left panel shows DN-POLG and right panel depicts data for the NDI1/AOX (N=2 each cell type). (B) Screen shot of a gene exemplifying DEGs that follow the pattern of cluster B in DN-POLG cells (see Fig. 1B). Top represents gene structure, the peak on the left falls on the annotated promoter; peak on the right is on the gene body in a non-annotated area. (C) Depiction of the percentage of peaks identified in the DN-POLG cells based on genomic locations. The odds ratio of a differentially expressed gene (DEG) having a change in H3K9ac occupancy was calculated based on the expected vs observed instances in which it occurred in DEGs and in non-DEGS.

**Fig. S2.**
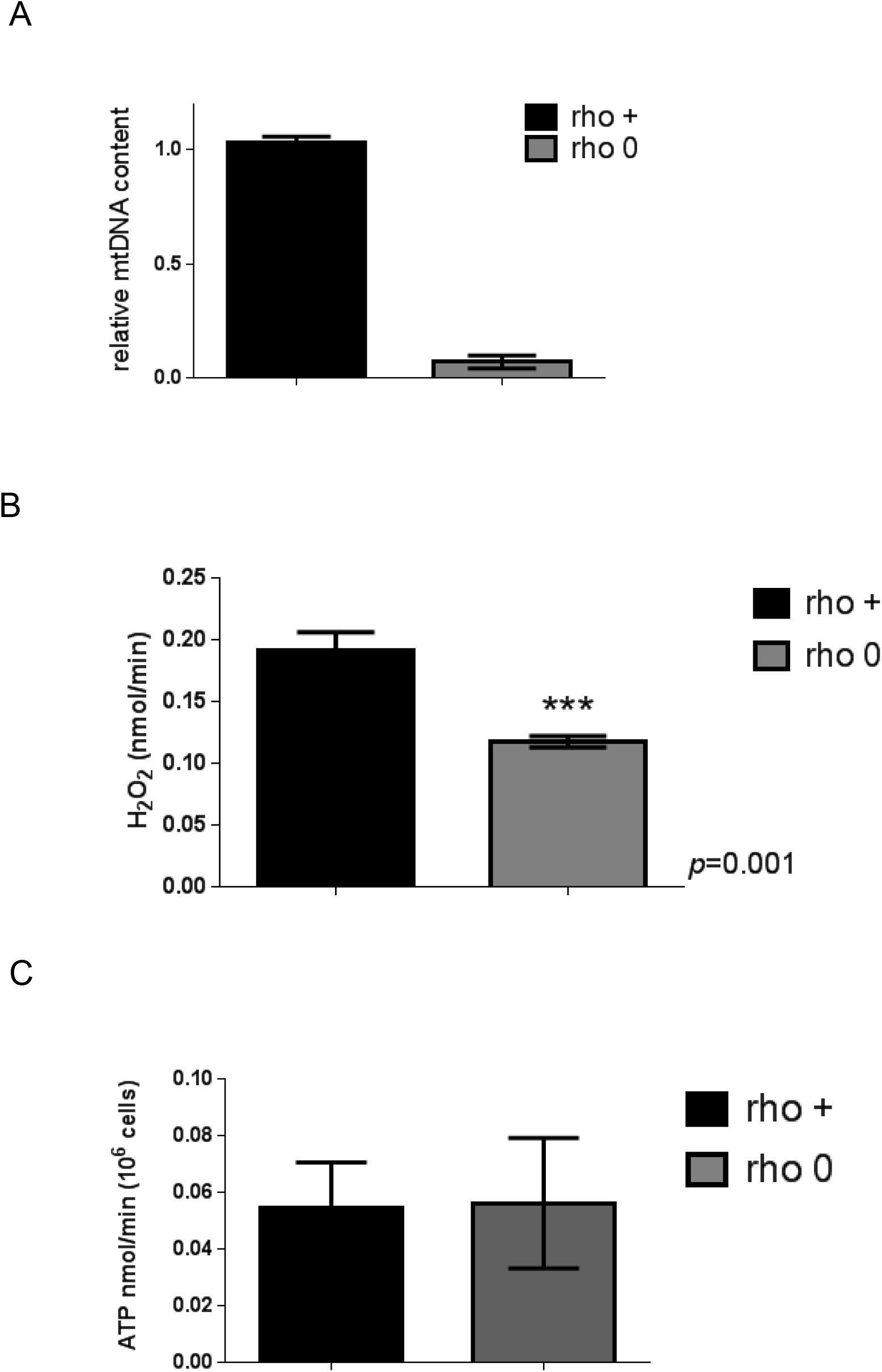
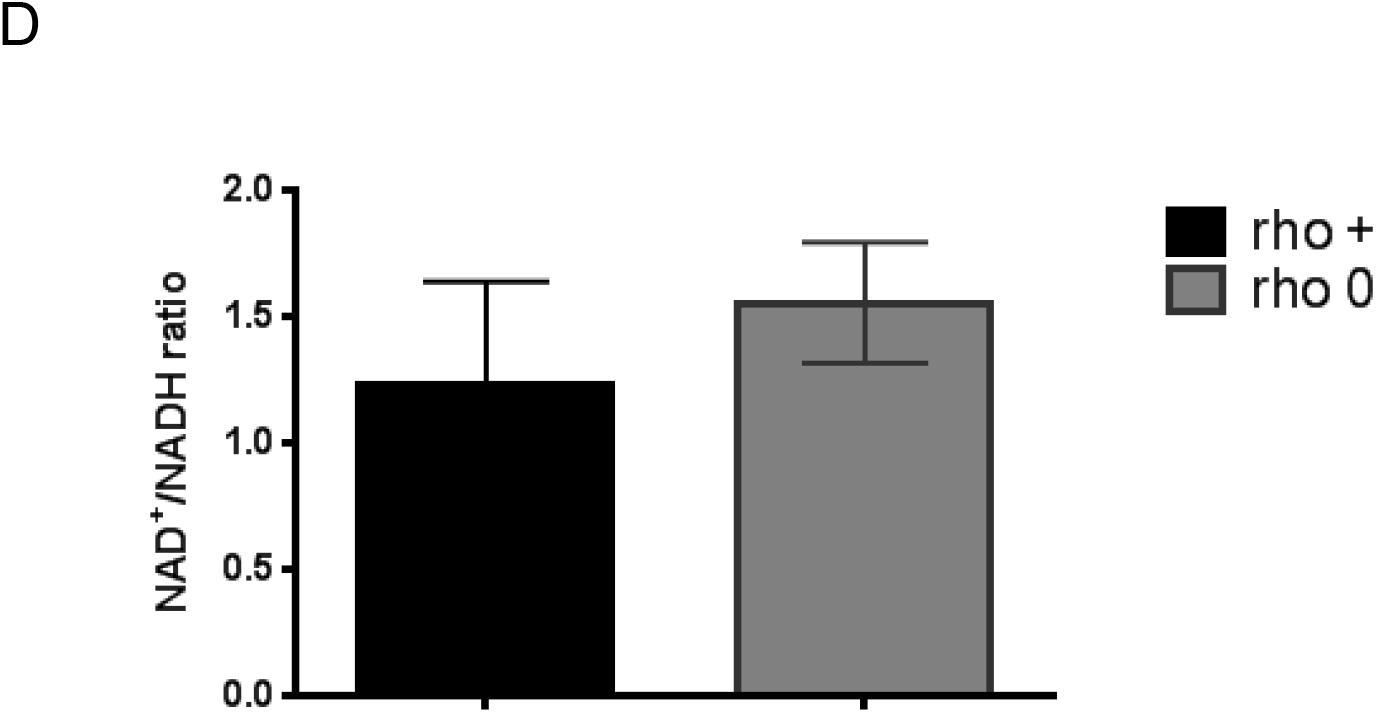
Biochemical parameters of 143B cells. (A) Relative mtDNA content was estimated using quantitative PCR; N=3; error bars depict SEM. (B) Levels of H_2_O_2_ were determined using Amplex red and a standard H_2_O_2_ curve. Data were normalized to cell number; N=3. (C) ATP levels were estimated using a fluorescent kit and deproteinized cellular lysates. N=3; data were normalized to cell number. (D) NAD^+^/NADH ratios were estimated using a commercially available kit in which the NAD^+^ or NADH-containing pool is measured separately. The ratio between the two is depicted in the graph. N=3.

**Fig. S3.**
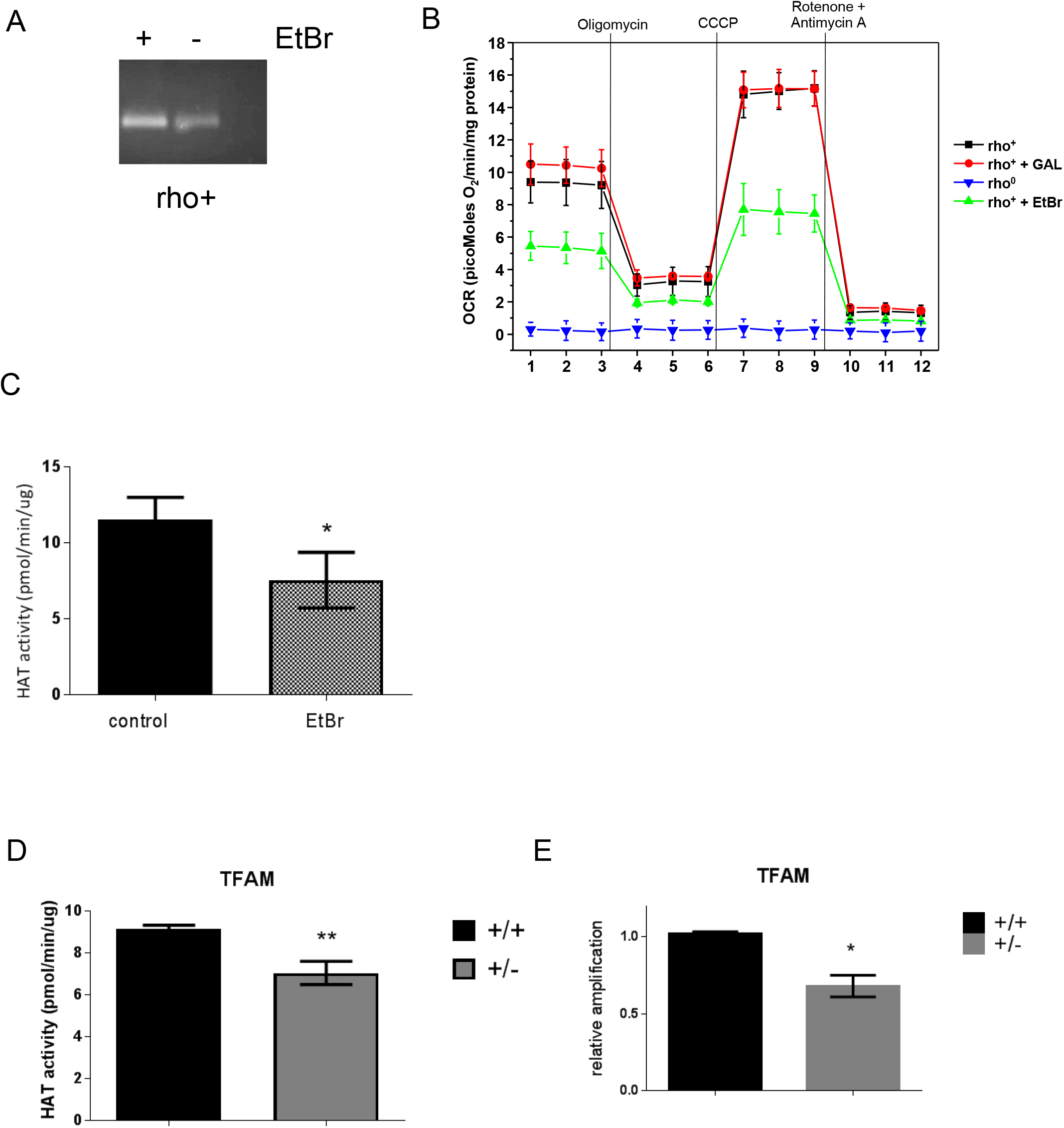
Decreased levels of mtDNA do not support full HAT activity. (A) PCR products from a 221bp fragment of the mtDNA in rho+ before and after treatment with EtBr for 1 week was ran on a 2% agarose gel. Reactions included the same amount of starting genomic DNA (15 ng) for 20 cycles. (B) Bioenergetics parameters of the cells was determined by extracellular flux analysis using the SeaHorse. Individual traces for oxygen consumption rate, OCR, are depicted upon addition of the various ETC inhibitors, including olygomycin, rotenone, antimycin and the uncoupler CCCP. Data were normalized to protein concentration; each sample was present in quadruplicates. Samples included: control (rho+) grown under glucose or galactose supplementation, rho+freshly treated with ethidium bromide for 7 days and rho0. (C) HAT activity was performed in rho+ cells treated with EtBr for 2 weeks (N=3) and (D) using TFAM wild type or heterozygote MEFs (N=3 per cell type). (E) Relative mtDNA content in the same samples was estimated as in A relative to TFAM +/+.

**Fig. S4.**
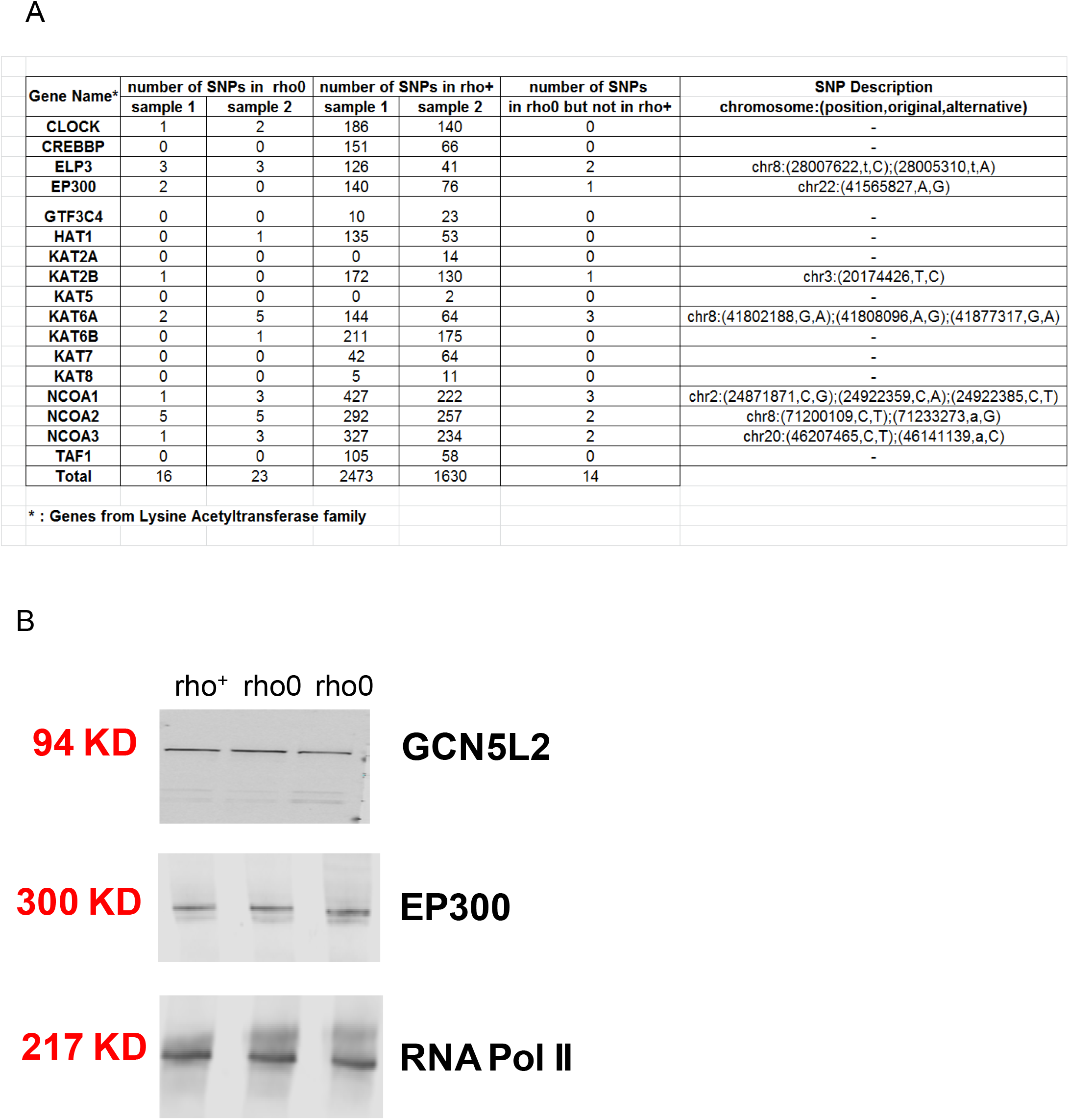
Changes in mutation rate or protein abundance are not observed in HATs when comparing rho+ to rho0 cells. (A) The sequence of the listed genes obtained based on deep-sequencing of two independent samples of input DNA was queried against the reference HG19 genome. The number of base changes (single nucleotide polymorphisms, SNPs) identified relative to the sequence contained in the reference HG19 genome is shown for each sample. The number of SNPs identified in the rho0 samples that were not identified in the rho+ control is also shown. For those, the genomic location and the exact base change is depicted (last column). (B) Western blots probing levels of the two main HATs that acetylate H3K9 and/or H3K27 was performed in whole cell lysates of rho+ or rho0 cells. RNA pol II was used as loading control.

**Fig. S5.**
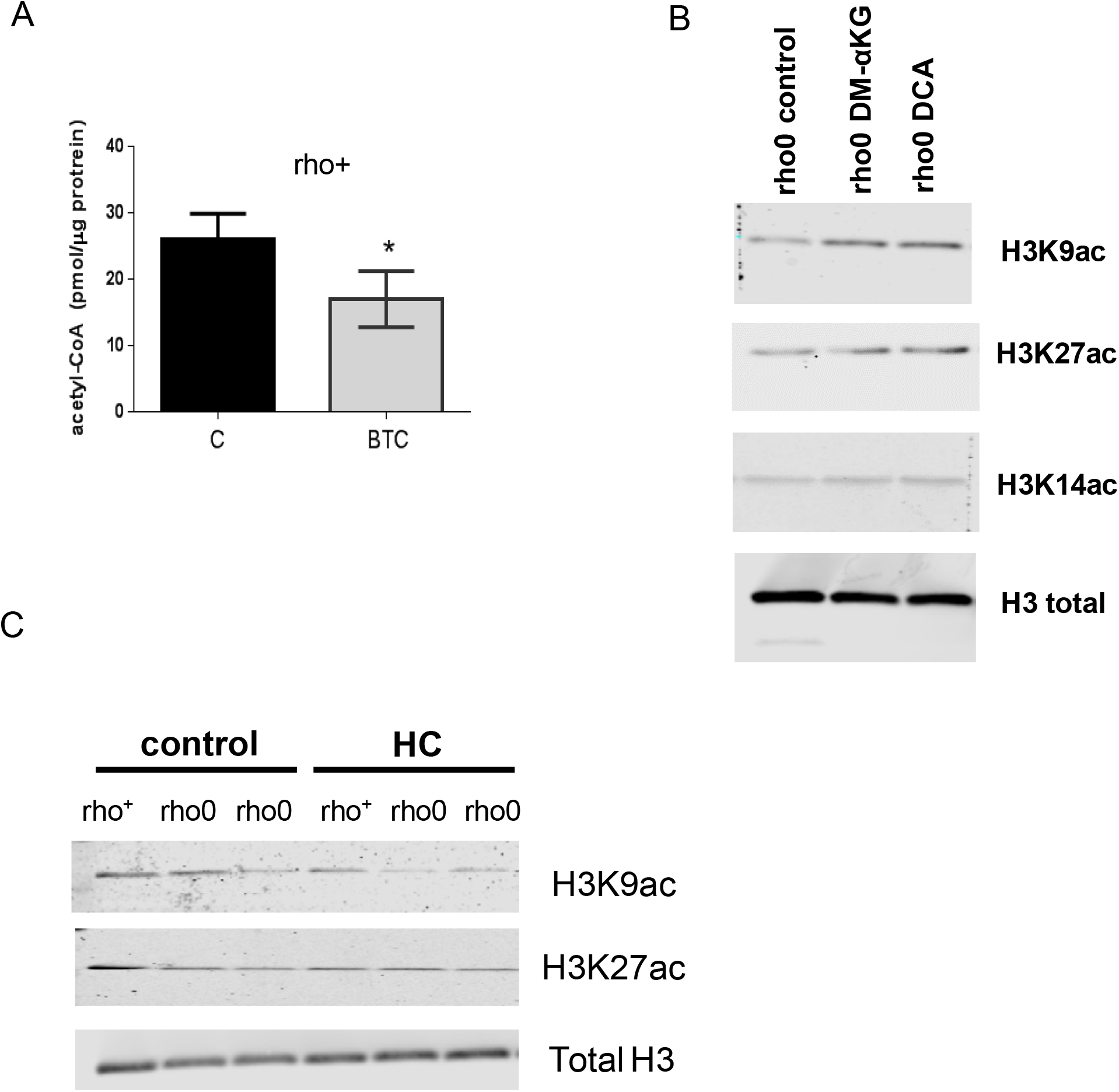
Changes in acetyl-CoA levels and in histone acetylation are detected after manipulation of the mitochondrial pool of acetyl-CoA in 143B cells. (A) 143B rho+ cells were exposed for 4h to BTC, a mitochondrial citrate carrier inhibitor. Total levels of acetyl-CoA were then estimated using deproteinized samples with a fluoresncence-based assay and an acetyl-CoA standard curve; N=3. (B). Histones were extracted and Western blots probing different histone acetylation marks performed after treatment of rho0 cells for 4h with DM-α-KG or DCA; Westerns are representative. (C) Representative Western blots probing different acetylated histone residues upon treatment of rho+ or rho0 cells with HC, an inhibitor of the cytosolic ATP citrate lyase.

**Fig. S6.**
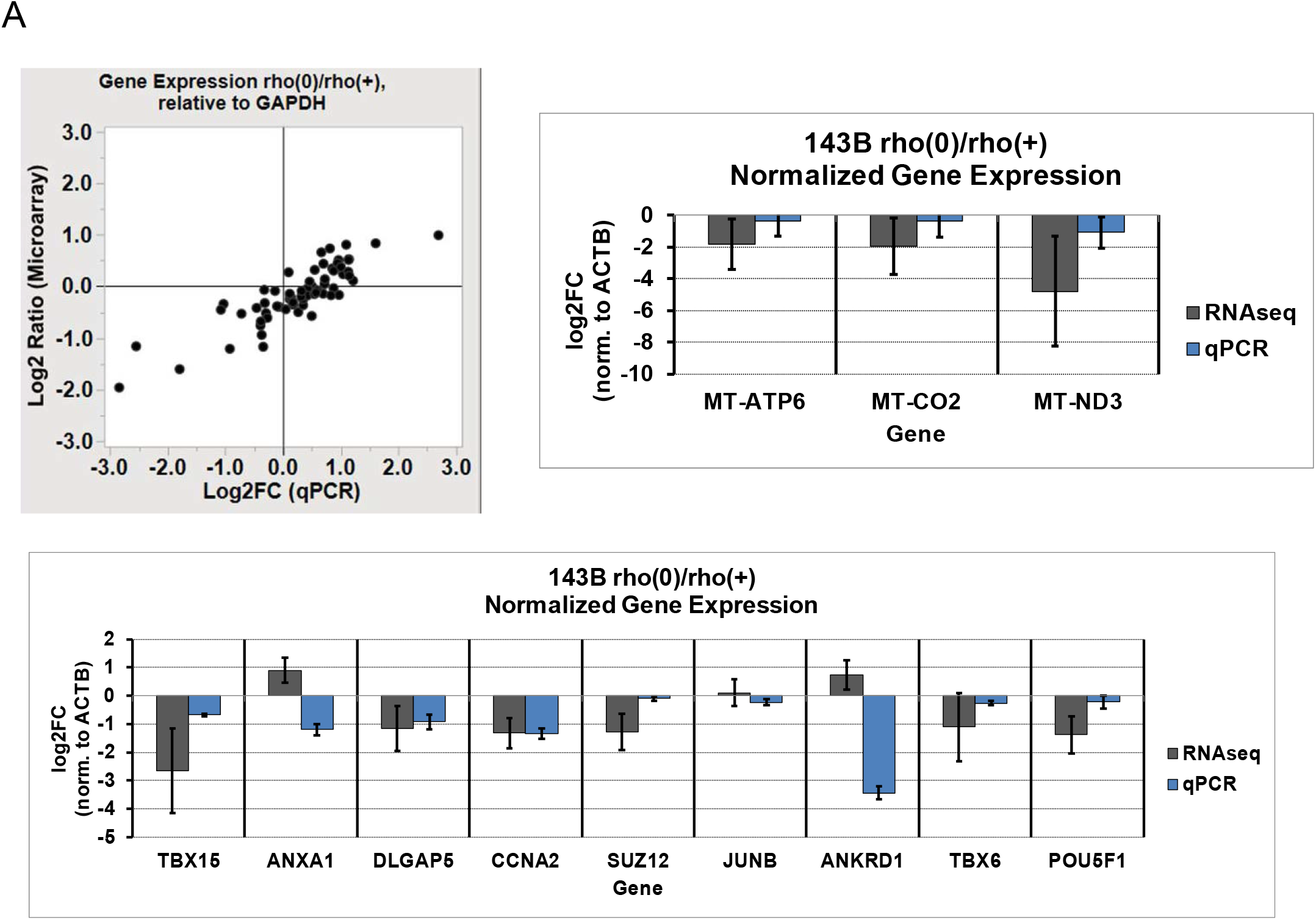
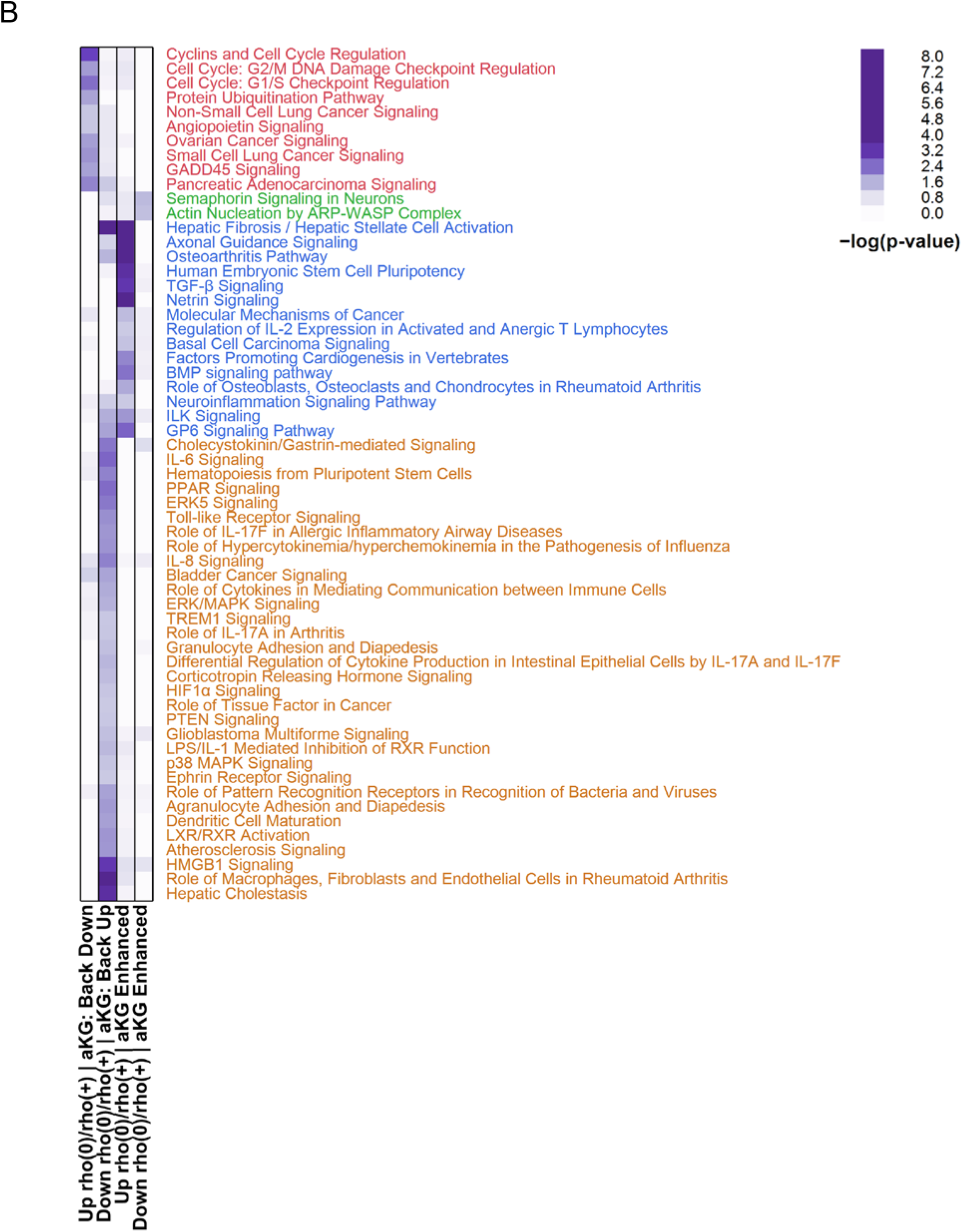
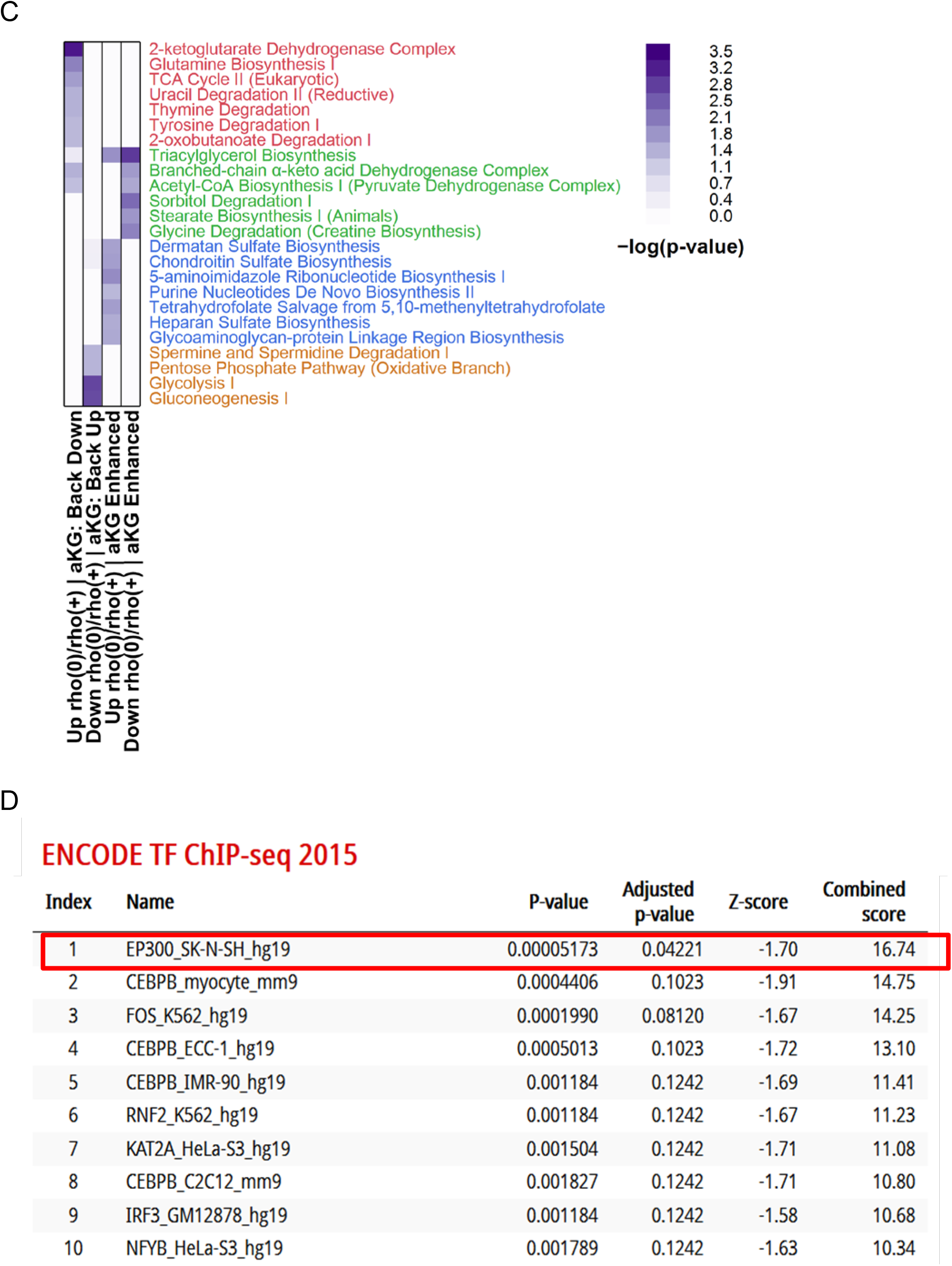
Validation and pathway enrichment analysis of genes rescued by DM-α-KG treatment of rho0 cells based on microarray experiments. (A) qPCR validation of genes detected in 143B cells by microarrays. Left upper panel depicts concordance of relative expression estimates in 143B rho0 v. rho+ cells normalized to house-keeping glyceraldehyde 3-phosphate dehydrogenase (GAPDH) between microarray (HG-U133 Plus 2.0, Affymetrix) and qPCR experiments (Human Amino Acid Metabolism Tier I PrimePCR panel, Biorad) for 65 nuclear-encoded genes; qPCR was performed from cDNA templates derived using the same total RNA extracts for both techniques (N=3 per cell derivative), and were performed in technical triplicates. Right upper and bottom panels show graphs of relative expression estimates in 143B rho0 v. rho+ cells normalized to ß-actin (ACTB) between RNA-seq and qPCR experiments (Custom PrimePCR panel, Biorad) for 3 mtDNA-encoded genes (right upper panel) and 9 nuclear DNA-encoded genes (bottom panel); qPCR was performed from cDNA templates derived using the same total RNA extracts for both techniques (N=3 per cell derivative), and were performed in technical triplicates. The genes identified by microarrays in the rho0 cells relative to rho+ whose expression were affected by treatment with DM-α-KG were used to define enriched pathways based on IPA analysis. (B) canonical pathways (C) only metabolic pathways. (D) The list of genes whose expression was reversed by DM-α-KG supplementation was used to retrieve ENCODE-based ChIP-seq information. The table indicate the proteins against which the ChIP-seq experiments were performed, followed by cell type and reference genome (Hg19, human, mm9, mouse).

**Fig. S7.**
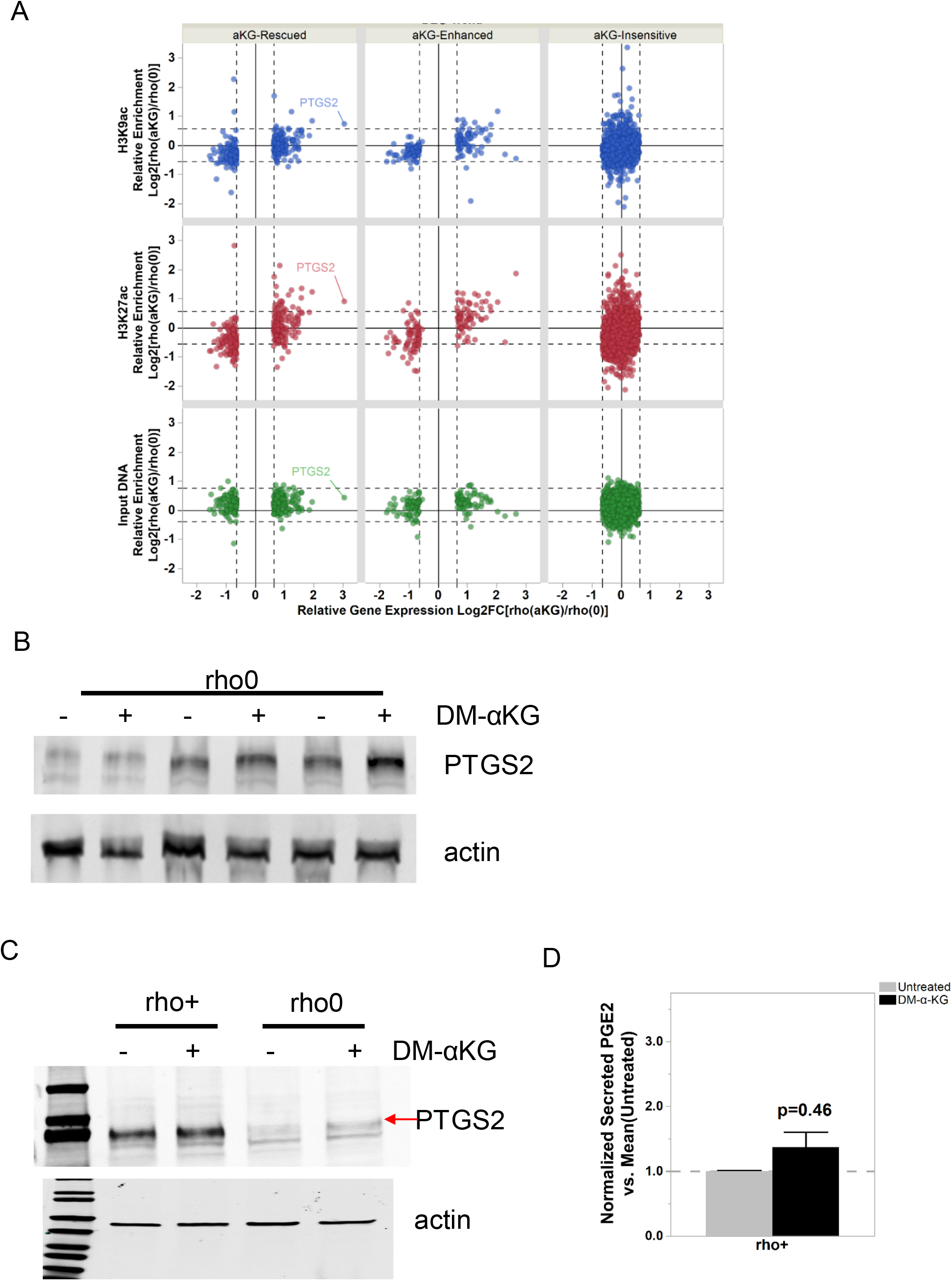
Effects of DM-α-KG treatment on histone marks and prostaglandin levels. Concordance between gene expression directionality (x-axis) and relative enrichment levels (y-axis) of H3K9ac (blue), H3K27ac (red) and input DNA (green) ChIPseq read densities within 2 Kb around the transcription start site (TSS) of genes responsive to supplementation with 20 mM DM-α-KG in 143B rho0 before [rho(0)] and after [rho(aKG)] 4-hr treatment. Each dot represents a gene; PTGS2 is highlighted as a reference. (B) Western blots for PTGS2 in 3 independent biological replicates of rho0 cells treated with DM-α-KG (including from Fig. 4E). (C) Same as (B) but when rho+ cells and rho0 were exposed to DM-α-KG for 4h in parallel experiments. (D) Levels of PGE2 were estimated by ELISA in the supernatant of the rho+ cells used in (C) prior to and after 4h exposure to DM-α-KG; N=3. Data was normalized to total protein content.

